# Chemolithoautotrophic acidophile phenotyping by measuring metabolic byproduct formation in microplates

**DOI:** 10.64898/2026.11.20.768213

**Authors:** Daniel A. Dailey, Mallori A. Herishko, Wei Wang, Lauren N. Murray, Sofia T. Bavaresco, Scott Banta

## Abstract

Chemolithoautotrophic acidophiles can support several low-pH biotechnologies, but their phenotyping remains limited by throughput and inconsistent methods. This study used an assay to monitor metabolic byproduct formation using absorbance indicators in static, oxic cultures in 96-well microplates. In abiotic experiments, ferric iron and/or pH measurements from 51 microplates containing simulant solutions span total iron concentrations of 0–1 M, pH 0.3–7.0, and incubation temperatures of 4–75 ◦C. Of these, 30 microplates tested different UV-transparency and gas-permeable membrane configurations. Another 12 microplates tested indicator compatibility with growth media containing ferrous iron and (in)soluble sulfur substrates. Biological experiments measured 438 microplates containing *Acidithiobacillus ferrooxidans* ATCC 23270, *Acidithiobacillus thiooxidans* ATCC 19377, or *Leptospirillum ferriphilum* DSM 14647 with chemical additives across 1,400 concentration conditions, yielding 142,038 time-course points, including abiotic controls. Measurements of apparent substrate oxidation and/or growth from 96 culture tubes and 99 shake flasks provide context for microplate observations. The miniaturized assay and this dataset may be useful for microbiology, (critical-mineral) biotechnology development, and synthetic biology.

## Background & Summary

The development of biotechnologies (for extreme operating conditions) is constrained by the physiological and metabolic properties of the host. Chemolithoautotrophic acidophiles span bacterial and archaeal lineages that perform carbon dioxide assimilation and obtain energy by oxidizing inorganic substrates.^1,2^ Oxidation of ferrous iron (Fe^2+^) and reduced sulfur (S) compounds, including elemental sulfur (S^0^), can regenerate ferric iron (Fe^3+^) and produce sulfuric acid (H_2_SO_4_), respectively, promoting metal dissolution from susceptible metal-bearing resources (e.g., rocks,^3^ plants^4^). Tolerance of acidity and certain dissolved metals enables their sustained use for metal extraction from sulfide resources.^5^ Laboratory studies also suggest potential for producing hydrocarbon biofuels^6^ and biomaterials, including biopolymers such as polyhydroxybutyrate^7^ and biogenic magnetic nanoparticles.^8^ Some members grow under aerobic or anaerobic conditions, and/or oxidize hydrogen for energy.^9,10^ This metabolic flexibility, together with genetic tools developed for selected bacteria^11,12^ and archaea,^13^ broadens possibilities for engineering low-pH bioprocesses. The potential applications of (engineered) chemolithoautotrophy may even extend beyond Earth.^14,15^

Predictable use of these non-model microorganisms nevertheless requires better characterization of strain-level physiology. Some members of *Acidithiobacillus* are well studied.^16^ Even so, their physiological characterization has lagged behind genome sequencing,^17^ and isolates exhibit substantial phenotypic variation.^18^ Revised species classifications further complicate comparison across studies.^17,19^ Phenotypes also depend on interacting factors including pH,^20,21^ redox potential,^21^ metal speciation and complexation,^22^ electron-donor identity,^16,23^ oxygen transfer,^24^ and, if solids are present, poorly understood interfacial properties.^21,23^ Testing combinations of strains, substrates, and cultivation conditions is therefore needed. However, limited high-throughput cultivation and phenotypic screening capabilities constrain the Test stage of Design–Build– Test–Learn (DBTL) workflows.^25^

Miniaturized assays can increase cultivation throughput^26,27^ and reveal condition-specific phenotypes or functions of poorly characterized genes.^28^ Standardized measurements^29,30^ could support DBTL workflows combining automated experimentation^31^ with data-driven experiment selection,^32^ perhaps within a self-driving laboratory.^33^ However, optical biomass measurements commonly used for model organisms are challenged in acidophilic cultures by low cell densities^34^ and interference from Fe^3+^, insoluble S, and precipitates.^35^ Biomass accumulation can decouple from substrate oxidation,^36,37^ which is often the function required for biotechnological applications. Monitoring apparent Fe^3+^ generation^38,39^ and acidification associated with S oxidation^4,23,40^ provides an alternative, requiring medium-specific calibration and condition-matched abiotic controls.

In this Data Descriptor, we present time-resolved absorbance measurements^41^ from a miniaturized assay^42^ in static, oxic cultures of chemolithoautotrophic acidophilic bacteria in 96-well microplates.^27^ The assay monitored two metabolic byproducts. The absorbance at 380 nm indicated apparent Fe^3+^ generation during Fe^2+^ oxidation,^39^ and the thymolsulfonephthalein (thymol blue) ratio indicated acidification associated with H^+^ release during sulfur oxidation.^40,43,44^ Feasibility and extendability tests validate these absorbance indicators in simulant solutions across total Fe concentrations of 0–1 M, pH 0.3–7.0, and incubation temperatures of 4–75 *^◦^*C within different microplate configurations (51 microplates; Tables 1–2). Compatibility tests evaluated the detectability of these absorbance indicators in acidophilic growth media containing Fe^2+^, thiosulfate (S_2_O_3_^2−^), tetrathionate (S_4_O_6_^2−^), brown or yellow S^0^, and selected substrate combinations (12 microplates; Table 3). Data from 88 demonstration tests monitored the apparent metabolism of *A. ferrooxidans* ATCC 23270, *A. thiooxidans* ATCC 19377, or *L. ferriphilum* DSM 14647 under exposure to 1,400 tested concentration conditions, including baseline medium controls (438 microplates; 142,038 time-course points; Table 4). The tested concentration conditions spanned about 10*^−^*^1^^1^–10^0^ M ranges for 3–58 added chemicals for each acidophile. Data from additional validation tests performed in shake flasks and culture tubes are provided for comparison of some of the results from this new miniaturized assay to those obtained using conventional experimental methods (Tables 5–6). These data have not previously been presented in another research article. These data can support evaluation of absorbance indicator versatility, comparisons of condition-dependent responses to conventional assays, and development of standardized acidophile phenotyping assays.

**Table 1:**
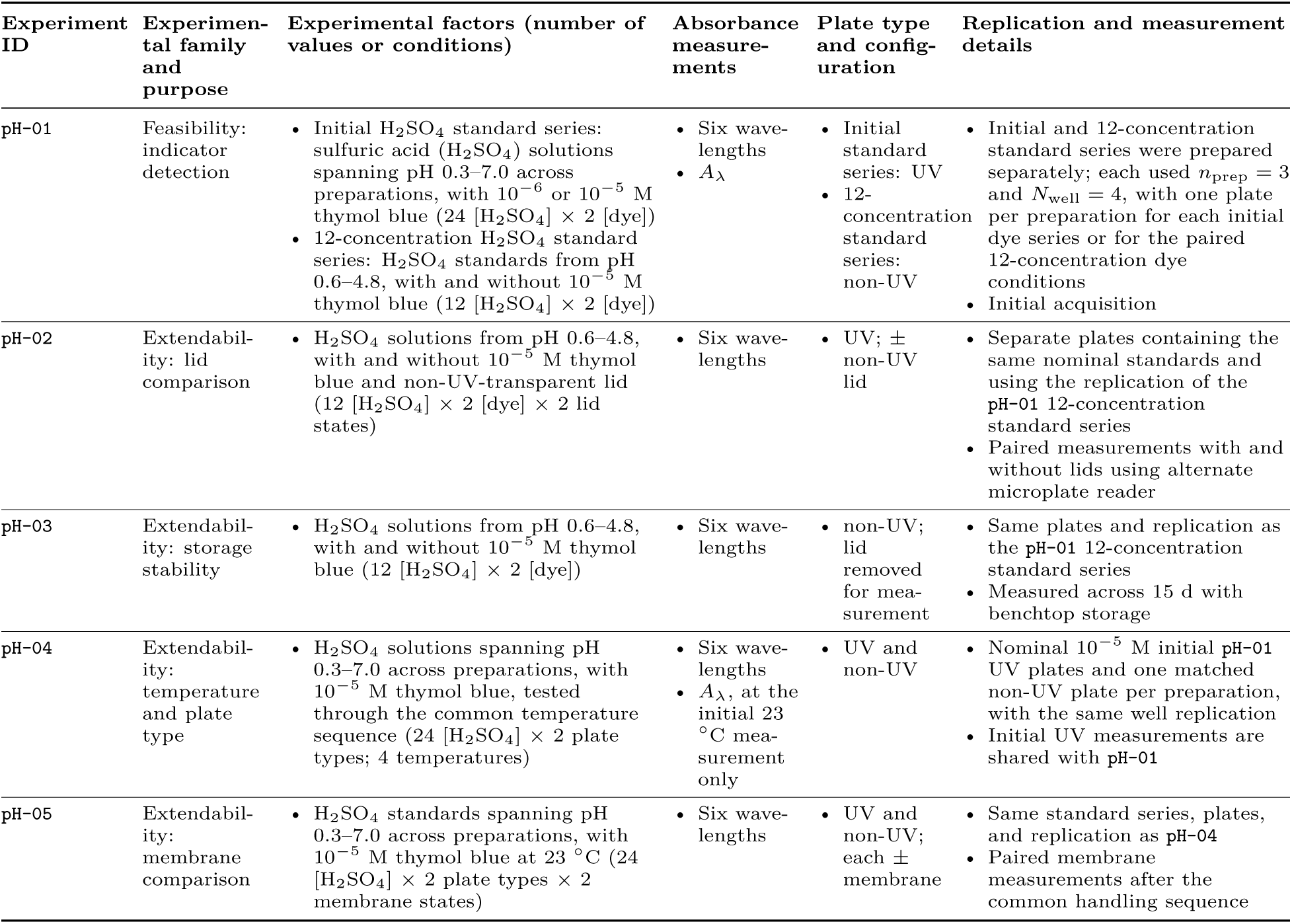
Feasibility and extendability tests of the thymol blue ratiometric pH indicator. Spectral measurements are denoted *A_λ_*, where *λ* spans 200–700 nm for the first preparation of each initial calibration series and 200–750 nm for the remaining scans, in 10 nm increments.

| Experiment ID | Experimen-tal family and purpose | Experimental factors (number of values or conditions) | Absorbance measure-ments | Plate type and config-uration | Replication and measurement details |
|---|---|---|---|---|---|
| pH-01 | Feasibility: indicator detection | • Initial H<sub>2</sub>SO<sub>4</sub> standard series: sulfuric acid (H<sub>2</sub>SO<sub>4</sub>) solutions spanning pH 0.3–7.0 across preparations, with 10<sup>−6</sup> or 10<sup>−5</sup> M thymol blue (24 [H<sub>2</sub>SO<sub>4</sub>] × 2 [dye])<br>• 12-concentration H<sub>2</sub>SO<sub>4</sub> standard series: H<sub>2</sub>SO<sub>4</sub> standards from pH 0.6–4.8, with and without 10<sup>−5</sup> M thymol blue (12 [H<sub>2</sub>SO<sub>4</sub>] × 2 [dye]) | • Six wave-lengths<br>• *A*<sub>λ</sub> | • Initial standard series: UV<br>• 12-concentration standard series: non-UV | • Initial and 12-concentration standard series were prepared separately; each used *n*<sub>prep</sub> = 3 and *N*<sub>well</sub> = 4, with one plate per preparation for each initial dye series or for the paired 12-concentration dye conditions<br>• Initial acquisition |
| pH-02 | Extendabil-ity: lid comparison | • H<sub>2</sub>SO<sub>4</sub> solutions from pH 0.6–4.8, with and without 10<sup>−5</sup> M thymol blue and non-UV-transparent lid (12 [H<sub>2</sub>SO<sub>4</sub>] × 2 [dye] × 2 lid states) | • Six wave-lengths | • UV; ± non-UV lid | • Separate plates containing the same nominal standards and using the replication of the pH-01 12-concentration standard series<br>• Paired measurements with and without lids using alternate microplate reader |
| pH-03 | Extendabil-ity: storage stability | • H<sub>2</sub>SO<sub>4</sub> solutions from pH 0.6–4.8, with and without 10<sup>−5</sup> M thymol blue (12 [H<sub>2</sub>SO<sub>4</sub>] × 2 [dye]) | • Six wave-lengths | • non-UV; lid removed for mea-surement | • Same plates and replication as the pH-01 12-concentration standard series<br>• Measured across 15 d with benchtop storage |
| pH-04 | Extendabil-ity: temperature and plate type | • H<sub>2</sub>SO<sub>4</sub> solutions spanning pH 0.3–7.0 across preparations, with 10<sup>−5</sup> M thymol blue, tested through the common temperature sequence (24 [H<sub>2</sub>SO<sub>4</sub>] × 2 plate types; 4 temperatures) | • Six wave-lengths<br>• *A*<sub>λ</sub>, at the initial 23 °C mea-surement only | • UV and non-UV | • Nominal 10<sup>−5</sup> M initial pH-01 UV plates and one matched non-UV plate per preparation, with the same well replication<br>• Initial UV measurements are shared with pH-01 |
| pH-05 | Extendabil-ity: membrane comparison | • H<sub>2</sub>SO<sub>4</sub> standards spanning pH 0.3–7.0 across preparations, with 10<sup>−5</sup> M thymol blue at 23 °C (24 [H<sub>2</sub>SO<sub>4</sub>] × 2 plate types × 2 membrane states) | • Six wave-lengths | • UV and non-UV; each ± membrane | • Same standard series, plates, and replication as pH-04<br>• Paired membrane measurements after the common handling sequence |

**Table 2:**
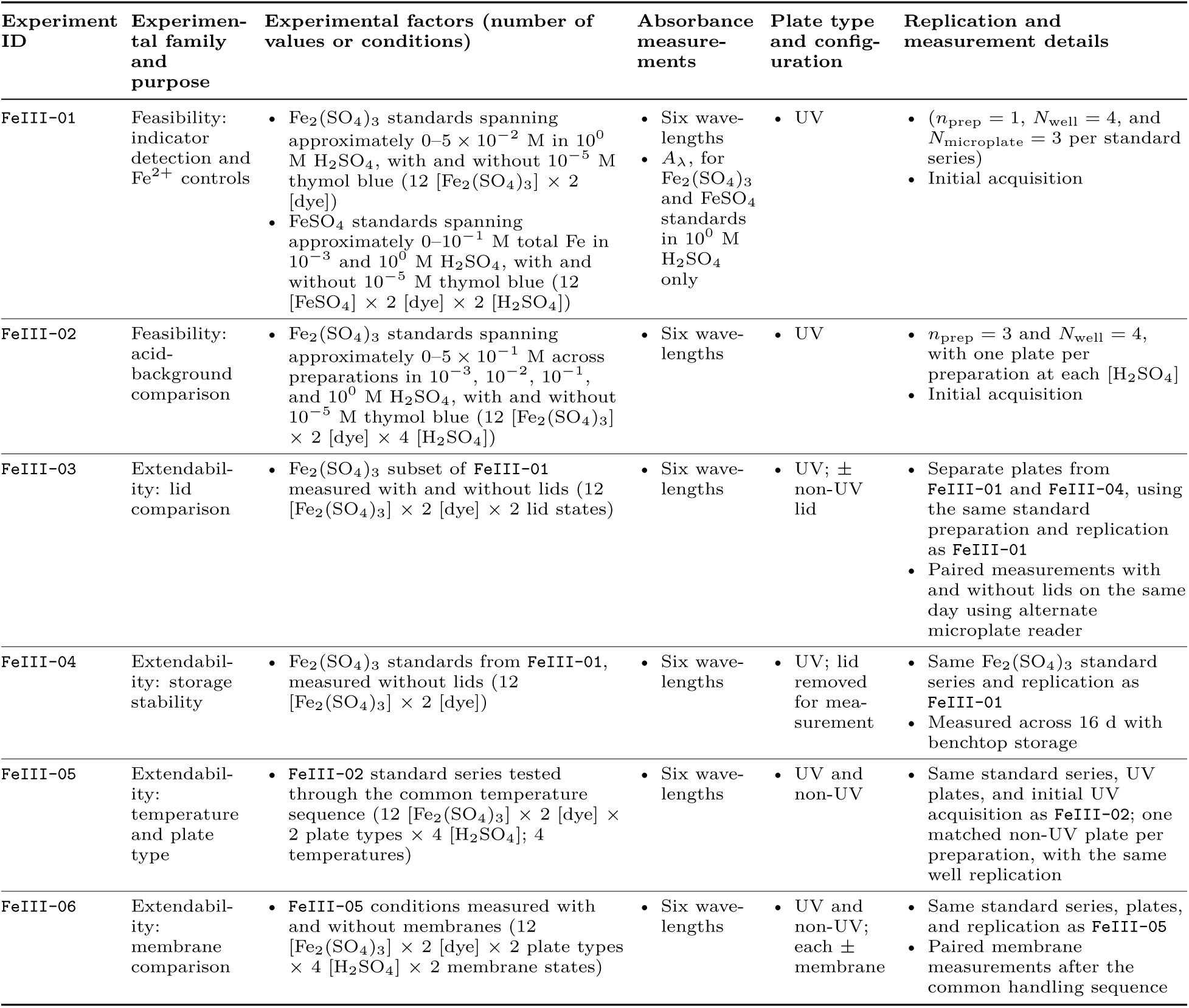
Feasibility and extendability tests of the Fe^3+^ absorbance indicator. Concentration ranges describe the supplying compound; Fe_2_(SO_4_)_3_ concentrations were calculated from measured total Fe using the stated stoichiometry. Both salt and acid concentrations refer to the standards before dye addition. Spectral measurements are denoted *A_λ_*, where *λ* spans 200–750 nm in 10 nm increments.

| Experiment ID | Experimen-tal family and purpose | Experimental factors (number of values or conditions) | Absorbance measure-ments | Plate type and config-uration | Replication and measurement details |
|---|---|---|---|---|---|
| FeIII-01 | Feasibility: indicator detection and Fe<sup>2+</sup> controls | • Fe<sub>2</sub>(SO<sub>4</sub>)<sub>3</sub> standards spanning approximately 0–5 × 10<sup>−2</sup> M in 10<sup>0</sup> M H<sub>2</sub>SO<sub>4</sub>, with and without 10<sup>−5</sup> M thymol blue (12 [Fe<sub>2</sub>(SO<sub>4</sub>)<sub>3</sub>] × 2 [dye])<br>• FeSO<sub>4</sub> standards spanning approximately 0–10<sup>−1</sup> M total Fe in 10<sup>−3</sup> and 10<sup>0</sup> M H<sub>2</sub>SO<sub>4</sub>, with and without 10<sup>−5</sup> M thymol blue (12 [FeSO<sub>4</sub>] × 2 [dye] × 2 [H<sub>2</sub>SO<sub>4</sub>]) | • Six wave-lengths<br>• *A*<sub>λ</sub>, for Fe<sub>2</sub>(SO<sub>4</sub>)<sub>3</sub> and FeSO<sub>4</sub> standards in 10<sup>0</sup> M H<sub>2</sub>SO<sub>4</sub> only | • UV | • (*n*<sub>prep</sub> = 1, *N*<sub>well</sub> = 4, and *N*<sub>microplate</sub> = 3 per standard series)<br>• Initial acquisition |
| FeIII-02 | Feasibility: acid-background comparison | • Fe<sub>2</sub>(SO<sub>4</sub>)<sub>3</sub> standards spanning approximately 0–5 × 10<sup>−1</sup> M across preparations in 10<sup>−3</sup>, 10<sup>−2</sup>, 10<sup>−1</sup>, and 10<sup>0</sup> M H<sub>2</sub>SO<sub>4</sub>, with and without 10<sup>−5</sup> M thymol blue (12 [Fe<sub>2</sub>(SO<sub>4</sub>)<sub>3</sub>] × 2 [dye] × 4 [H<sub>2</sub>SO<sub>4</sub>]) | • Six wave-lengths | • UV | • *n*<sub>prep</sub> = 3 and *N*<sub>well</sub> = 4, with one plate per preparation at each [H<sub>2</sub>SO<sub>4</sub>]<br>• Initial acquisition |
| FeIII-03 | Extendabil-ity: lid comparison | • Fe<sub>2</sub>(SO<sub>4</sub>)<sub>3</sub> subset of FeIII-01 measured with and without lids (12 [Fe<sub>2</sub>(SO<sub>4</sub>)<sub>3</sub>] × 2 [dye] × 2 lid states) | • Six wave-lengths | • UV; ± non-UV lid | • Separate plates from FeIII-01 and FeIII-04, using the same standard preparation and replication as FeIII-01<br>• Paired measurements with and without lids on the same day using alternate microplate reader |
| FeIII-04 | Extendabil-ity: storage stability | • Fe<sub>2</sub>(SO<sub>4</sub>)<sub>3</sub> standards from FeIII-01, measured without lids (12 [Fe<sub>2</sub>(SO<sub>4</sub>)<sub>3</sub>] × 2 [dye]) | • Six wave-lengths | • UV; lid removed for mea-surement | • Same Fe<sub>2</sub>(SO<sub>4</sub>)<sub>3</sub> standard series and replication as FeIII-01<br>• Measured across 16 d with benchtop storage |
| FeIII-05 | Extendabil-ity: temperature and plate type | • FeIII-02 standard series tested through the common temperature sequence (12 [Fe<sub>2</sub>(SO<sub>4</sub>)<sub>3</sub>] × 2 [dye] × 2 plate types × 4 [H<sub>2</sub>SO<sub>4</sub>]; 4 temperatures) | • Six wave-lengths | • UV and non-UV | • Same standard series, UV plates, and initial UV acquisition as FeIII-02; one matched non-UV plate per preparation, with the same well replication |
| FeIII-06 | Extendabil-ity: membrane comparison | • FeIII-05 conditions measured with and without membranes (12 [Fe<sub>2</sub>(SO<sub>4</sub>)<sub>3</sub>] × 2 [dye] × 2 plate types × 4 [H<sub>2</sub>SO<sub>4</sub>] × 2 membrane states) | • Six wave-lengths | • UV and non-UV; each ± membrane | • Same standard series, plates, and replication as FeIII-05<br>• Paired membrane measurements after the common handling sequence |

**Table 3:** Compatibility tests of absorbance indicators in chemolithoautotrophic acidophile media. Added Fe_2_(SO_4_)_3_ concentrations exclude the Fe already present in the medium. Measured pH values describe medium–standard mixtures before dye addition. Medium-component concentrations refer to the standard medium before dye addition. All measurements were abiotic six-wavelength absorbance measurements in non-UV-transparent microplates without lids.

| Experiment ID | Experimental family and purpose | Experimental factors (number of values or conditions) | Replication and measurement details |
|---|---|---|---|
| AFM-01 | Compatibility: Fe<sup>3+</sup> indicator | • AFM1 and AFM3 with 0–5.2 × 10<sup>−2</sup> M added Fe<sub>2</sub>(SO<sub>4</sub>)<sub>3</sub>, without dye (8 [Fe<sub>2</sub>(SO<sub>4</sub>)<sub>3</sub>] × 2 media) | • AFM1 and AFM3 occupy separate well groups on the same plates |
| F2S-01 | Compatibility: ratiometric pH and Fe<sup>3+</sup> indicators | • F2S with 0–5.2 × 10<sup>−2</sup> M added Fe<sub>2</sub>(SO<sub>4</sub>)<sub>3</sub>, without dye (8 [Fe<sub>2</sub>(SO<sub>4</sub>)<sub>3</sub>])<br>• F2S supplemented with H<sub>2</sub>SO<sub>4</sub>, spanning measured pH 0.7–2.1, with and without 10<sup>−5</sup> M thymol blue (8 [H<sub>2</sub>SO<sub>4</sub>] × 2 [dye]) | • Fe<sub>2</sub>(SO<sub>4</sub>)<sub>3</sub> and H<sub>2</sub>SO<sub>4</sub> standard series occupy separate well groups on the same plates |
| Starkey-01 | Compatibility: ratiometric pH indicator | • Starkey (St) medium with brown or yellow S<sup>0</sup>, supplemented with H<sub>2</sub>SO<sub>4</sub>, spanning measured pH 0.7–2.8 for each medium, with and without 10<sup>−5</sup> M thymol blue (8 [H<sub>2</sub>SO<sub>4</sub>] × 2 S<sup>0</sup> types × 2 [dye]) | • Both S<sup>0</sup> types occupy separate well groups on the same plates |
| 9K-01 | Compatibility: ratiometric pH indicator | • AFM4 or AFM5 supplemented with H<sub>2</sub>SO<sub>4</sub>, spanning measured pH 0.7–3.8 for each medium, with and without 10<sup>−5</sup> M thymol blue (8 [H<sub>2</sub>SO<sub>4</sub>] × 2 media × 2 [dye]) | • AFM4 and AFM5 occupy separate well groups on the same plates |

**Table 4:**
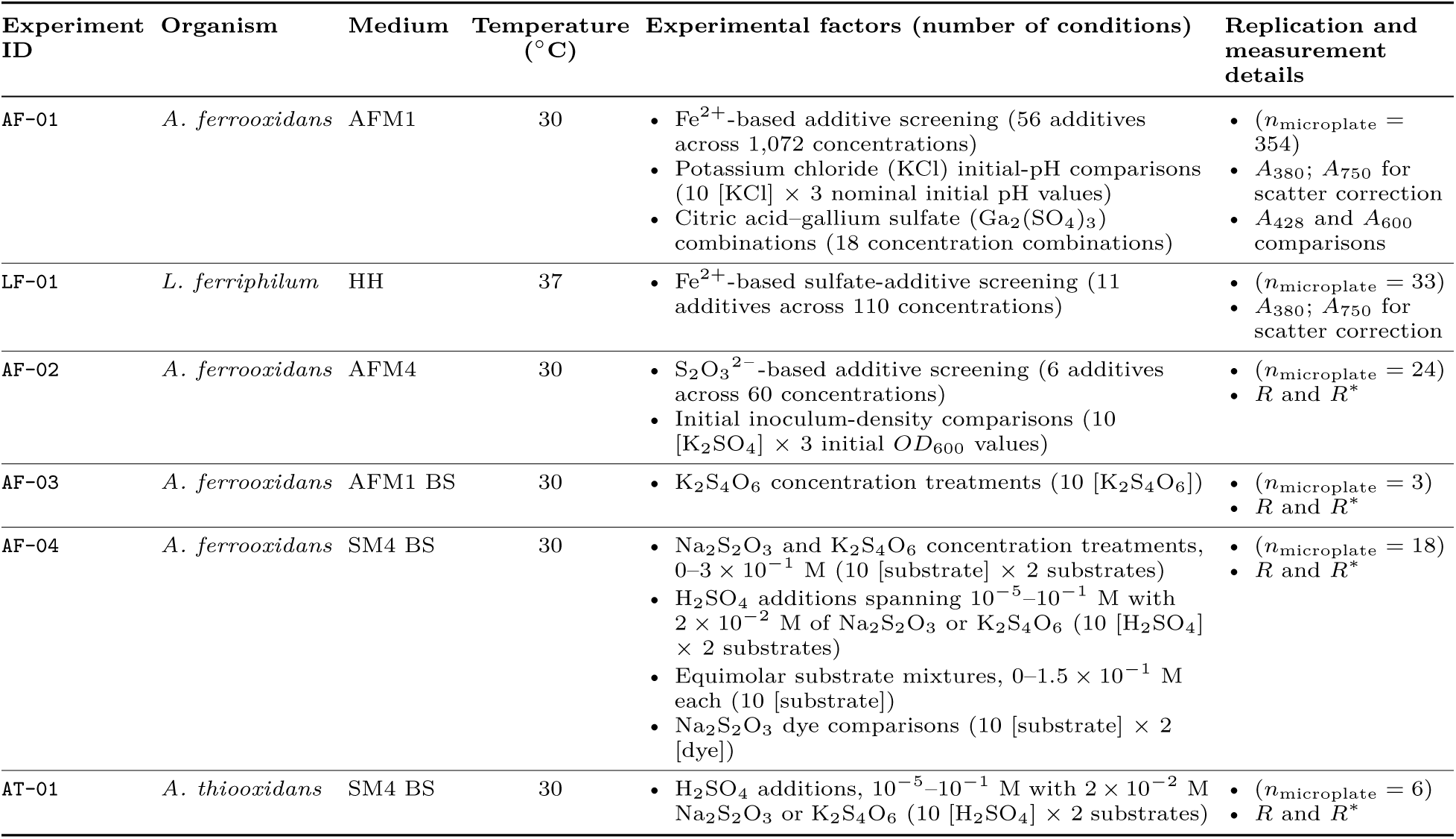
Demonstration tests in acidophile microplate cultures. The six experiment groups comprise independently inoculated microplates (*n*_microplate_ = 438 in total). *A*_380_ indicates apparent formation of the metabolic byproduct Fe^3+^ in cultures supplied with Fe^2+^. Measured *R* and scatter-corrected *R^∗^* indicate acidification associated with metabolic H^+^ release in soluble-S cultures, as defined in Data processing. Additive concentrations use the deposited compound basis, except explicitly specified elemental-Ga concentrations in citric-acid/Ga_2_(SO_4_)_3_ combinations.

| Experiment ID | Organism | Medium | Temperature (°C) | Experimental factors (number of conditions) | Replication and measurement details |
|---|---|---|---|---|---|
| AF-01 | *A. ferrooxidans* | AFM1 | 30 | • Fe<sup>2+</sup>-based additive screening (56 additives across 1,072 concentrations)<br>• Potassium chloride (KCl) initial-pH comparisons (10 [KCl] × 3 nominal initial pH values)<br>• Citric acid–gallium sulfate (Ga<sub>2</sub>(SO<sub>4</sub>)<sub>3</sub>) combinations (18 concentration combinations) | • (*n*<sub>microplate</sub> = 354)<br>• *A*<sub>380</sub>; *A*<sub>750</sub> for scatter correction<br>• *A*<sub>428</sub> and *A*<sub>600</sub> comparisons |
| LF-01 | *L. ferriphilum* | HH | 37 | • Fe<sup>2+</sup>-based sulfate-additive screening (11 additives across 110 concentrations) | • (*n*<sub>microplate</sub> = 33)<br>• *A*<sub>380</sub>; *A*<sub>750</sub> for scatter correction |
| AF-02 | *A. ferrooxidans* | AFM4 | 30 | • S<sub>2</sub>O<sub>3</sub><sup>2−</sup>-based additive screening (6 additives across 60 concentrations)<br>• Initial inoculum-density comparisons (10 [K<sub>2</sub>SO<sub>4</sub>] × 3 initial *OD*<sub>600</sub> values) | • (*n*<sub>microplate</sub> = 24)<br>• *R* and *R*<sup>\*</sup> |
| AF-03 | *A. ferrooxidans* | AFM1 BS | 30 | • K<sub>2</sub>S<sub>4</sub>O<sub>6</sub> concentration treatments (10 [K<sub>2</sub>S<sub>4</sub>O<sub>6</sub>]) | • (*n*<sub>microplate</sub> = 3)<br>• *R* and *R*<sup>\*</sup> |
| AF-04 | *A. ferrooxidans* | SM4 BS | 30 | • Na<sub>2</sub>S<sub>2</sub>O<sub>3</sub> and K<sub>2</sub>S<sub>4</sub>O<sub>6</sub> concentration treatments, 0–3 × 10<sup>−1</sup> M (10 [substrate] × 2 substrates)<br>• H<sub>2</sub>SO<sub>4</sub> additions spanning 10<sup>−5</sup>–10<sup>−1</sup> M with 2 × 10<sup>−2</sup> M of Na<sub>2</sub>S<sub>2</sub>O<sub>3</sub> or K<sub>2</sub>S<sub>4</sub>O<sub>6</sub> (10 [H<sub>2</sub>SO<sub>4</sub>] × 2 substrates)<br>• Equimolar substrate mixtures, 0–1.5 × 10<sup>−1</sup> M each (10 [substrate])<br>• Na<sub>2</sub>S<sub>2</sub>O<sub>3</sub> dye comparisons (10 [substrate] × 2 [dye]) | • (*n*<sub>microplate</sub> = 18)<br>• *R* and *R*<sup>\*</sup> |
| AT-01 | *A. thiooxidans* | SM4 BS | 30 | • H<sub>2</sub>SO<sub>4</sub> additions, 10<sup>−5</sup>–10<sup>−1</sup> M with 2 × 10<sup>−2</sup> M Na<sub>2</sub>S<sub>2</sub>O<sub>3</sub> or K<sub>2</sub>S<sub>4</sub>O<sub>6</sub> (10 [H<sub>2</sub>SO<sub>4</sub>] × 2 substrates) | • (*n*<sub>microplate</sub> = 6)<br>• *R* and *R*<sup>\*</sup> |

**Table 5:** Validation tests in acidophile shake-flask cultures using AFM1 medium. All conditions include pH, ORP, and cerimetry measurements. Fluorescence denotes SYBR Green I measurements. Nominal final metal-salt treatment concentrations are expressed on a cation basis; labels identify the supplying compound.

| Experiment ID | Organism | Temperature (°C) | Experimental factors (number of conditions) | Replication and measurement details |
|---|---|---|---|---|
| FL-01 | *A. ferrooxidans* | 30 | • Biotic: 0–10<sup>−1</sup> M Ga<sub>2</sub>(SO<sub>4</sub>)<sub>3</sub> (7 [Ga<sub>2</sub>(SO<sub>4</sub>)<sub>3</sub>])<br>• Abiotic: 0–10<sup>−1</sup> M Ga<sub>2</sub>(SO<sub>4</sub>)<sub>3</sub> (3 [Ga<sub>2</sub>(SO<sub>4</sub>)<sub>3</sub>]) | • Soluble Fe and Ga at four sample-collection times<br>• (Non-)planktonic fluorescence |
| FL-02 | *A. ferrooxidans* | 30 | • Biotic: no-additive control; 10<sup>−1</sup> M Ga<sub>2</sub>(SO<sub>4</sub>)<sub>3</sub>; 10<sup>−1</sup> M CuSO<sub>4</sub>; or 5 × 10<sup>−2</sup> M NaCl (4 conditions) | • Planktonic fluorescence<br>• Initial soluble Fe: all conditions; Ga and Cu: corresponding metal-addition conditions |
| FL-03 | *A. ferrooxidans* | 30 | • Biotic: no-additive control; 5 × 10<sup>−2</sup> M citric acid; 10<sup>−1</sup> M Ga<sub>2</sub>(SO<sub>4</sub>)<sub>3</sub>; 10<sup>−1</sup> M VOSO<sub>4</sub>; or 10<sup>−1</sup> M Al<sub>2</sub>(SO<sub>4</sub>)<sub>3</sub> (5 conditions) | • Planktonic fluorescence<br>• Final non-planktonic fluorescence in the no-additive, Ga<sub>2</sub>(SO<sub>4</sub>)<sub>3</sub>, VOSO<sub>4</sub>, and Al<sub>2</sub>(SO<sub>4</sub>)<sub>3</sub> conditions<br>• Initial soluble Fe: the same four conditions; Ga, V, and Al: corresponding metal-addition conditions |
| FL-04 | *A. ferrooxidans* | 30 | • Biotic: 0 or 10<sup>−3</sup> M Ga<sub>2</sub>(SO<sub>4</sub>)<sub>3</sub>, each with 0, 10<sup>−3</sup> or 6 × 10<sup>−3</sup> M citric acid (2 [Ga<sub>2</sub>(SO<sub>4</sub>)<sub>3</sub>] × 3 [citric acid])<br>• Abiotic: 10<sup>−3</sup> M Ga<sub>2</sub>(SO<sub>4</sub>)<sub>3</sub> (1 condition) | • (Non-)planktonic fluorescence<br>• Initial soluble Ga |
| FL-05 | *A. ferrooxidans* | 30 | • Biotic: 0 or 10<sup>−3</sup> M Ga<sub>2</sub>(SO<sub>4</sub>)<sub>3</sub>, each with 0, 1.2 × 10<sup>−2</sup> or 1.8 × 10<sup>−2</sup> M citric acid (2 [Ga<sub>2</sub>(SO<sub>4</sub>)<sub>3</sub>] × 3 [citric acid])<br>• Abiotic: no-additive control (1 condition) | • Planktonic fluorescence<br>• Non-planktonic fluorescence in the biotic no-additive control condition<br>• Initial soluble Ga |

**Table 6:** Validation tests in acidophile culture tubes. TB-02 includes ORP time courses, and TB-03– TB-04 include pH time courses; all include planktonic SYBR Green I fluorescence at the final acquisition. TB-01 used the original microplate inocula for visual assessment. Treatment concentrations are nominal values before inoculation; metal-salt concentrations are expressed on a cation basis, and energy-substrate concentrations refer to the supplying compounds.

| Experiment ID | Organism | Medium | Temperature (°C) | Experimental factors (number of conditions) |
|---|---|---|---|---|
| TB-01 | • Abiotic<br>• *L. ferriphilum* | HH | 37 | • No-additive control condition |
| TB-02 | • Abiotic<br>• *A. ferrooxidans*<br>• *A. ferridurans*<br>• *L. ferriphilum* | HH | 37 | • No-additive control; 10<sup>−1</sup> M Ga<sub>2</sub>(SO<sub>4</sub>)<sub>3</sub>; or 10<sup>−2</sup> M In<sub>2</sub>(SO<sub>4</sub>)<sub>3</sub> (4 organism/control groups × 3 conditions) |
| TB-03 | • Abiotic<br>• *A. ferrooxidans*<br>• *A. thiooxidans* | SM4 BS | 30 | • Brown S<sup>0</sup> (1 g/L) with 0 or 10<sup>−1</sup> M Ga<sub>2</sub>(SO<sub>4</sub>)<sub>3</sub> (3 organism/control groups × 2 [Ga<sub>2</sub>(SO<sub>4</sub>)<sub>3</sub>]) |
| TB-03 | • Abiotic<br>• *A. caldus* | SM4 BS | 37 | • Brown S<sup>0</sup> (1 g/L) with 0 or 10<sup>−1</sup> M Ga<sub>2</sub>(SO<sub>4</sub>)<sub>3</sub> (2 organism/control groups × 2 [Ga<sub>2</sub>(SO<sub>4</sub>)<sub>3</sub>]) |
| TB-04 | • Abiotic<br>• *A. ferrooxidans*<br>• *A. thiooxidans* | SM4 BS | 30 | • K<sub>2</sub>S<sub>4</sub>O<sub>6</sub> (5 × 10<sup>−3</sup> M) with 0 or 10<sup>−1</sup> M Ga<sub>2</sub>(SO<sub>4</sub>)<sub>3</sub> (3 organism/control groups × 2 [Ga<sub>2</sub>(SO<sub>4</sub>)<sub>3</sub>]) |
| TB-04 | • Abiotic<br>• *A. caldus* | SM4 BS | 37 | • K<sub>2</sub>S<sub>4</sub>O<sub>6</sub> (5 × 10<sup>−3</sup> M) with 0 or 10<sup>−1</sup> M Ga<sub>2</sub>(SO<sub>4</sub>)<sub>3</sub> (2 organism/control groups × 2 [Ga<sub>2</sub>(SO<sub>4</sub>)<sub>3</sub>]) |

## Methods

### Materials

Ultrapure water (H_2_O) was used in all experiments unless otherwise noted. Chemicals, materials, equipment, bacterial strains, media formulations, and nominal or measured stock concentrations are described in the dataset.^41^ The term basal salts (BS) denotes the specified medium formulation without its added energy substrate.

### Preparation of additive stock solutions

Additive stock solutions were prepared in H_2_O and adjusted using H_2_SO_4_ and sodium hydroxide to pH 1.80 *±* 0.05, except for metal stocks used in AFM4 microplate experiments, which were adjusted to pH 3.00 *±* 0.05. Stock pH was measured using a pH 700 benchtop meter (Oakton) calibrated at least daily using pH 1.68, 4.01, and 7.00 standards. Additive stock solutions were prepared at 2*×* the target concentrations in the medium–additive mixture (200 µL) before dye addition. Nominal stock concentrations were calculated from reagent masses and final solution volumes, using 50 mL CertiTube™ digestion tubes (Environmental Express). Solubility information, where available, guided the selection of target stock concentrations. Soluble elemental concentrations in some additive stocks were measured by inductively coupled plasma optical emission spectroscopy (ICP-OES; Horiba Ultima Expert) as previously described.^4^ Treatment labels identify the added compound, unless otherwise noted. Conversion between compound and cation concentrations used the corresponding stoichiometry.

For each additive stock measured by ICP-OES, an aliquot of the freshly prepared 2*×* stock was diluted in the analytical matrix without prior centrifugation or filtration and used for comparison with the filtered aliquots in the solubility assessment. The analytical matrix contained 2% (w/v) HNO_3_; titanium and zirconium stocks instead used 2% (w/v) HCl. Separate stock aliquots (1 mL) were centrifuged (17,000 *×* g, 10 min) immediately after preparation and after 7 d of storage under ambient laboratory conditions. Additional stock aliquots (500 µL) were mixed with 2*×* AFM1 medium (500 µL) and centrifuged under the same conditions at both time points. Supernatants were passed through 0.2 µm filters and diluted in the corresponding analytical matrix for elemental analysis. Stock-only filtrates were additionally diluted twofold. For stock–AFM1 mixtures, the soluble elemental contribution from AFM1 was subtracted before comparison with the corresponding uncentrifuged, unfiltered stock aliquot. For stocks quantified by ICP-OES, the solubility-assessment acceptance criterion was an absolute relative difference of *≤* 3% from the corresponding uncentrifuged, unfiltered stock aliquot at both time points. For stocks assigned nominal concentrations, aliquots were centrifuged using the same stock-only procedure. The visual and pH stability acceptance criteria were no visible solids before or after centrifugation at either time point and an absolute pH change over 7 d of *≤* 0.05, respectively.

For both measured and nominal stocks, serial dilutions were prepared in H_2_O acidified to the corresponding stock pH using H_2_SO_4_, sterile-filtered, and stored alongside the 2*×* stocks in the dark at room temperature until use. Concentrations after dilution were calculated from the measured or nominal parent-stock concentration and the cumulative dilution factor. Concentrations calculated from measured stocks were distinguished from directly measured final concentrations. No individual dilution factor exceeded tenfold.

### Microplate preparation and absorbance acquisition

Independent starter cultures are counted as *n*_starter_; *n*_batch_ counts groups of cultures initiated together under the same preparation conditions. Technical biotic and abiotic wells, repeated measurements, and titrations are denoted by the corresponding *N* subscripts. For abiotic tests, *n*_prep_ denotes independent standard preparations, *N*_microplate_ denotes technical microplates receiving the same preparation, and *N*_well_ denotes technical wells per condition per microplate. For biological tests, *n*_microplate_, *n*_flask_, and *n*_tube_ denote separately inoculated vessels. Biological microplates are the independent replicate units, and their within-condition wells are technical replicates despite receiving separate inocula. Experiment-specific sample sizes are given below, with analysis-specific availability in Data processing. For microplate configurations, UV and non-UV denote UV-transparent and non-UV-transparent, respectively; *±* denotes measurements with and without the indicated lid or membrane. A thymol blue stock solution (2.0 *×* 10*^−^*^4^ M) was prepared in aqueous ethanol (10% (v/v)) adjusted to pH 4 using H_2_SO_4_. A portion was diluted tenfold in the same solution to obtain a 2.0 *×* 10*^−^*^5^ M stock for the initial pH calibration. Abiotic wells contained 200 µL of standard or medium–standard mixture, with or without an additional 10 µL of the dye stock. The two stocks gave calculated final dye concentrations of approximately 9.5 *×* 10*^−^*^6^ and 9.5 *×* 10*^−^*^7^ M, designated nominally as 10*^−^*^5^ and 10*^−^*^6^ M, respectively. Absorbance measurements were performed using a Molecular Devices SpectraMax M2 microplate reader unless otherwise noted. Six-wavelength acquisitions comprised 380,^39^ 428,^42^ 430,^44^ 485, 545,^44^ and 750 nm.^45^ Measurements were acquired without lids except during the paired lid comparison, which used a Tecan Infinite M Plex microplate reader and sequential acquisitions from the same uncovered and covered microplate on the same day.

### Feasibility and extendability tests of the ratiometric pH indicator

For the initial H_2_SO_4_ standard series (Table 1), the assigned dye stock was dispensed before the corresponding acid standard. Serial dilutions of 5 *×* 10*^−^*^1^ M H_2_SO_4_ were prepared in H_2_O, and dispensed using a multichannel pipette. These series were prepared independently on separate days, with deliberately varied measured pH values. The 12-concentration calibration, lid-comparison, and storage wells used the 2.0 *×* 10*^−^*^4^ M dye stock, with matched wells without dye. For the 12-concentration calibration, pH was measured after dye addition and assigned to the corresponding wells without dye. Microplates in the storage experiments were covered with low-evaporation lids between acquisitions; pH was not remeasured during storage.

Calibration measurements (pH-01) and the initial temperature-comparison acquisition (pH-04) followed incubation at 23 *^◦^*C for 30 min. For the temperature comparisons, the same microplates were subsequently incubated sequentially at 4, 30, and 75 *^◦^*C for 2 h each, with absorbance acquired immediately after each incubation. For membrane comparisons, microplates were then covered with lids, sealed with Parafilm, and incubated overnight at 23 *^◦^*C. Absorbance was acquired immediately before and after Breathe-Easy™ gas-exchange membranes were attached.

### Feasibility and extendability tests of the Fe^3+^ absorbance indicator

Fe_2_(SO_4_)_3_ standard series were made by serial dilution in the designated H_2_SO_4_ solutions; separate FeSO_4_ standard series served as Fe^2+^ control conditions (Table 2). Total Fe concentrations were measured by ICP-OES for each sample after serial dilution. For chemical measurements, aliquots (1 mL) were centrifuged (3,000 *×* g, 10 s), and supernatants (750 µL) were transferred to separate microcentrifuge tubes and centrifuged again (17,000 *×* g, 60 s). Supernatants were diluted in 2% (w/v) HNO_3_ for ICP-OES. The measured total Fe concentrations describe soluble Fe across oxidation states in the analysed supernatants after serial dilution and before dye addition. These chemical measurements were shared by the corresponding optical wells and did not represent separate chemical measurements of every well.

Final total Fe concentrations were calculated from the measured standard concentrations. For FeIII-02, the Fe_2_(SO_4_)_3_ standard series at each H_2_SO_4_ concentration were prepared independently on separate days. For FeIII-02, standard pH measurements were limited to the Fe_2_(SO_4_)_3_ standard series in 10*^−^*^3^ and 10*^−^*^2^ M H_2_SO_4_. For FeIII-01, cerimetry was performed on the Fe_2_(SO_4_)_3_ standards in 10^0^ M H_2_SO_4_ and on the FeSO_4_ standards in each of 10*^−^*^3^ and 10^0^ M H_2_SO_4_ (*N*_titration_ = 3 per standard). Clarified supernatant aliquots (250 µL) were mixed with H_2_O (750 µL) and ferroin indicator (10 µL), then titrated with standardized cerium(IV) sulfate (Ce(SO_4_)_2_) (10*^−^*^1^ M for the one-electron reaction, 1–10 µL increments) while vortexing until the indicator changed from red to blue as previously described.^4^

Initial acquisitions for FeIII-01, FeIII-02, and FeIII-05, subsequent temperature incubations, and membrane comparisons followed the pH-indicator procedures. Storage experiments used approximately 23°C, with low-evaporation lids removed for acquisition.

### Measuring absorbance indicators in chemolithoautotrophic acidophile media

To assess indicator compatibility with chemolithoautotrophic acidophile media (Table 3), media were prepared at 2*×* their standard concentrations in H_2_O. For pH characterization, 10^0^ M H_2_SO_4_ was serially diluted twofold in H_2_O. H_2_O replaced the acid standard in the zero-added-H_2_SO_4_ control condition. For characterization of the Fe^3+^ absorbance indicator, a 1.047 *×* 10*^−^*^1^ M Fe_2_(SO_4_)_3_ stock, calculated from the ICP-OES measurement of total Fe, was serially diluted twofold in 10^0^ M H_2_SO_4_.

Calibration solution (100 µL) was added to wells containing 2*×* medium (100 µL). Stocks containing insoluble S^0^ were vortexed vigorously before dispensing to improve the uniformity of the added S^0^ mass within the slurry. Only the pH-indicator experiments used the common dye-addition procedure, with matched wells without dye. The pH of the H_2_SO_4_ standards and of each medium–standard mixture was measured before dye addition. Measurements for each medium were shared across conditions with and without dye and across its technical microplates. Each experiment group in Table 3 used the shared calibration stocks with *n*_prep_ = 1, *N*_microplate_ = 3, and *N*_well_ = 3. Added Fe_2_(SO_4_)_3_ concentrations were calculated from the measured stock concentration, the compound stoichiometry, and the mixing volumes. All Fe_2_(SO_4_)_3_-addition conditions, including the zero-added-Fe control condition, received nominally 5 *×* 10*^−^*^1^ M H_2_SO_4_ from the calibration solution. Microplates were incubated at 23 *^◦^*C for 30 min before absorbance acquisition.

### Culturing and monitoring acidophiles in microplates

Experiments used *Acidithiobacillus ferrooxidans* ATCC 23270, *Acidithiobacillus thiooxidans* ATCC 19377, and *Leptospirillum ferriphilum* DSM 14647 (Table 4). Washed cells were suspended in AFM1 BS and normalized to *OD*_600_ = 5.0 *±* 0.1 (mean *±* SD, *N*_measurement_ = 3). The normalized suspension was added to 2*×* medium to give an *OD*_600_ twice the target initial value, then mixed with an equal volume of 2*×* additive solution (100 µL each) in non-UV-transparent microplates. The nominal initial *OD*_600_ before dye addition was 0.001, except for AFM4 additive experiments, which used 0.1; the AFM4 potassium sulfate (K_2_SO_4_) treatment conditions additionally included 0.01 and 0.001. For abiotic control conditions, an equivalent volume of sterile AFM1 BS replaced the cell suspension.

Each biological experiment used *n*_microplate_ = 3 per condition unless specified otherwise in Data processing. Each microplate used a separate set of inocula, with a different inoculum in each biotic well and condition-matched abiotic wells prepared alongside them (*N*_biotic well_ = *N*_abiotic well_ = 3 per condition). The same inocula supplied corresponding treatment and control conditions within a microplate, so those conditions remained paired. Different treatment conditions were analysed separately and were not counted as replicates of one another. Abiotic control conditions received the same medium, additives, and dye addition, but individual abiotic wells were not paired with individual inocula.

Fe^3+^-indicator wells contained no thymol blue and had a final volume of 200 µL. For ratiometric pH-indicator experiments, the thymol blue stock (10 µL, 2.0*×*10*^−^*^4^ M) was added to the 200 µL mixture, giving a nominal final dye concentration of 10*^−^*^5^ M. An additional *A. ferrooxidans* S_2_O_3_^2−^ experiment in SM4 BS used 30 µL of the same stock. Treatment concentrations in Table 4 describe the 200 µL medium–additive mixture before dye addition; final concentrations were calculated from the nominal or measured stock concentration, as applicable. Concentrations of compounds supplying S energy substrates refer to the supplying compound. Added Fe concentrations describe the additional Fe supplied by the treatment, excluding Fe already present in the medium. The initial SM4 pH was 3.95 *±* 0.03 (mean *±* SD, *N*_measurement_ = 3).

To reduce evaporation, sterile H_2_O (300 µL) was added to peripheral wells, low-evaporation lids were sealed with Parafilm between acquisitions, and open beakers of sterile H_2_O (100 mL) were kept beside the microplates. Water was not replenished in culture wells. Cultures were incubated under static, oxic conditions. Absorbance was acquired repeatedly from the same microplates without lids. The wavelengths used for each acquisition are specified in the instrument exports. Elapsed time was calculated from the initial acquisition for each microplate.

### Culturing and monitoring acidophiles in shake flasks

Validation cultures (100 mL) were established in 250 mL shake flasks as previously described^4^ (Table 5). Biotic shake flasks were inoculated to a nominal initial *OD*_600_ of 0.001 using approximately 20–50 µL of cell suspension, with the same inoculum volume within each experimental batch. Abiotic control conditions received the same volume of sterile medium. Each biotic or abiotic condition used *n*_flask_ = 3 per batch; biotic flasks received independent starter cultures, with corresponding starters reused across conditions. Shake flasks were incubated at 150 rpm at the indicated temperatures. Concentrations of sodium chloride (NaCl) and citric acid refer to the compounds added. Each biotic experimental batch included a no-additive control condition.

Samples (1 mL) were collected from the same shake flasks at successive time points for measurement. The pH was measured as described above. The oxidation–reduction potential (ORP) was analogously measured relative to a silver/silver chloride (Ag/AgCl) reference electrode with 3 *×* 10^0^ M potassium chloride (KCl) electrolyte. Fe^2+^ was again determined by cerimetry. The vanadium oxysulfate (VOSO_4_) treatment interfered with cerimetry,^46^ so the affected results were excluded from quantitative Fe^2+^ determination. Soluble Fe, gallium (Ga), copper (Cu), vanadium (V), and aluminum (Al) concentrations were measured by ICP-OES with radial viewing. (Non-)planktonic fraction recovery and SYBR Green I fluorescence measurements were performed as previously described.^4^ Initial pH, ORP, Fe^2+^ concentrations determined by cerimetry, and soluble elemental concentrations were assigned from measurements of the shared starting medium.

### Culturing and monitoring acidophiles in culture tubes

Additional strains used in culture-tube validation tests were *Acidithiobacillus ferridurans* DSM 29468 and *Acidithiobacillus caldus* ATCC 51756 (Table 6). SM4 BS used in these culture tubes omitted citric acid. Abiotic control conditions matched the treatment conditions and incubation temperatures.

For experiments TB-02–TB-04, equal volumes of 2*×* medium and 2*×* treatment solution were mixed to give 6 mL in 14 mL polystyrene culture tubes with dual-position caps. The treatment solutions had a target pH of 1.8; the SM4 BS stock had a target pH of 2.2. Concentrations in Table 6 are nominal values before inoculation. Independent starter cultures were established from freshly thawed cells, washed, and normalized. Each biotic condition received one tube per starter (*n*_starter_ = *n*_tube_ = 3 per organism), with the same starters reused across treatment conditions and a matched number of abiotic tubes. Each biotic culture tube received 20 µL of inoculum to give a nominal initial *OD*_600_ of 0.003; abiotic control conditions received an equivalent volume of cell-free BS. All culture tubes were incubated at 150 rpm with caps in the vented position and held at 60*^◦^* in mounted wire racks.

Cultures supplied with Fe^2+^ in HH medium were monitored by ORP for approximately 7 d, and cultures supplied with S substrates by pH for approximately 21 d, using the procedures described above. Evaporative losses were replaced with sterile ultrapure water before acquisition directly in the same culture tubes. Initial values were assigned from measurements of the shared medium–treatment mixture before inoculation. Final planktonic fluorescence measurements were performed.

Separately, the original *L. ferriphilum* inocula used in microplate experiments were added to HH medium without an added metal treatment in culture tubes when the microplate experiments ended (*n*_tube_ = 3). These culture tubes were incubated at 37 *^◦^*C and 150 rpm for 14 d and assessed visually for red coloration relative to the abiotic control condition. These inocula were distinct from the fresh starter cultures used in experiments TB-02–TB-04.

### Data processing

For the initial indicator feasibility tests, measured absorbance and derived ratios were evaluated for individual wells within each microplate before calculating summaries for each standard. Measured ratios use absorbance before scatter correction. The ratiometric pH indicator (*R*; unitless) was calculated using:

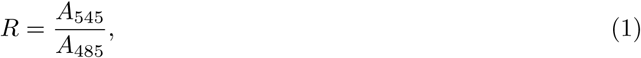

where *A*_545_ and *A*_485_ are the absorbances at 545 and 485 nm, respectively (AU).^44^ Scatter-corrected absorbance (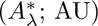; AU) was calculated using:

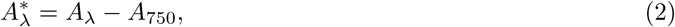

where *A_λ_*is the absorbance at wavelength *λ* and *A*_750_ is the absorbance at 750 nm (AU).^45^ The scatter-corrected ratiometric pH indicator (*R^∗^*) was calculated by substituting Equation (2) into Equation (1). Calculations excluded missing measurements, values that could not be interpreted as numbers, undefined or infinite values, and absorbance above the microplate reader limit of 4 AU. Ratios with denominators at or below 10*^−^*^6^ AU were excluded to avoid division by zero or values too close to zero. Negative finite absorbance or derived ratios were retained when measurements at the required wavelengths and the ratio denominator otherwise satisfied these criteria. Initial-calibration fits used the arithmetic mean for each standard only when all technical wells satisfied these criteria. Tests of mean paired differences or log-fold changes against zero used two-sided one-sample Student-*t* tests on the specified microplate or experimental-batch estimates. Unless stated otherwise, confidence intervals for means used the Student-*t* distribution and all intervals were pointwise 95% intervals. Holm adjustment was applied within the comparison families defined below.^47^ Optical comparison families included only tests that could be evaluated; unavailable tests had no adjusted probability. Biological comparison families retained all planned comparisons, including those that could not be evaluated.

For the ratiometric pH indicator, initial pH calibration curves were fitted to the mean *R*(∗) within each microplate as a function of the measured pH for each standard, using unweighted nonlinear least squares and a four-parameter logistic function^48^:

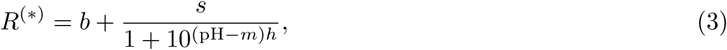

where *b* is the lower asymptote (unitless), *s* is the fitted response span (unitless), *m* is the midpoint pH (unitless), and *h* is the slope coefficient (unitless). Parameter and fitted-response uncertainties were estimated using HC3-adjusted^49^ linearized influence functions summed within each microplate. The covariance was adjusted for the number of microplates using:

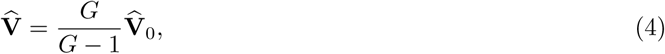

where **V̂**_0_ is the covariance calculated from the microplate-summed influence functions and *G* is the number of microplates. Approximate 95% confidence limits for each estimated parameter or fitted response were calculated using the Student-*t* distribution with *G −* 1 degrees of freedom.^50^ Model fits were assessed for convergence, numerical stability of the parameter estimates, and coverage of the measured standard range. Adjacent standards, ordered by measured pH or concentration, were compared using the signed difference between their means (*δX_j_*; AU or unitless):

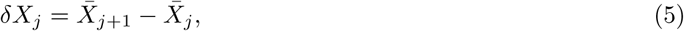

where *X̅_j_* and *s_j_* denote the arithmetic mean and sample standard deviation of well absorbance or derived ratios for standard *j*, respectively, with standards ordered by increasing measured pH or Fe concentration. Local sensitivity (*S_j_*; indicator units per pH unit or M) was calculated using:

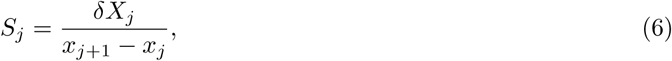

where *x_j_* is the measured pH or Fe concentration at standard *j*. The unsigned adjacent-standard separation score (*E_j_*; unitless) was calculated from Equation (5) using:

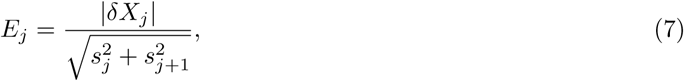

Differences opposite to the expected decrease with pH or increase with Fe concentration were classified as local reversals. Confidence intervals used Student-*t* limits for signed differences and sensitivities; positive separation scores were log-transformed before calculating intervals and back-transforming.

The observed response span (*W_X_* ; AU or unitless) within each microplate was calculated using:

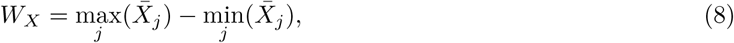

The pooled within-standard SD (*s*_pool_; AU or unitless) was calculated using:

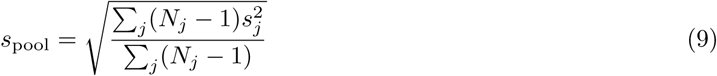

where *N_j_* is the number of eligible technical wells for standard *j*. The response-span-to-SD ratio (*Q*; unitless) was calculated using:

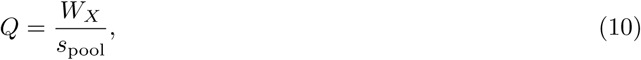

Measured and scatter-corrected values were paired using the same wells and standards. Log-fold changes (*L_M_* ; unitless) were calculated using:

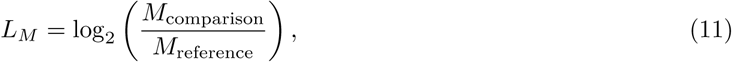

where the numerator and denominator are positive values of the same quantity for the comparison condition and its stated comparator, respectively. For scatter-correction comparisons, *M* was *W_X_*, *s*_pool_, or *Q*, with scatter-corrected and measured values as the numerator and denominator, respectively. Log ratios were calculated within each microplate and compared with zero within the comparison family comprising observed response span, pooled within-standard SD, and their ratio. Student-*t* intervals were back-transformed to express fold changes.

To assess pH prediction, each independently prepared H_2_SO_4_ standard series and its corresponding microplate were withheld in turn, and calibration curves fitted to the remaining preparations were used to predict pH in the withheld standards and in Fe_2_(SO_4_)_3_ mixtures.^51^ Comparisons used shared 0.25-pH bins with equal weight per bin. For each Fe_2_(SO_4_)_3_ mixture within a microplate, signed and squared pH prediction errors were averaged across the calibration fits before calculating bias and RMSE, respectively. Prediction bias (*B*_pH_; pH units) was calculated using:

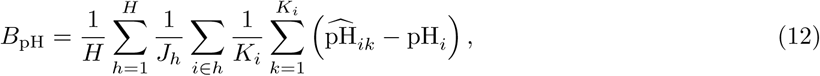

where *H* is the number of shared pH bins, *J_h_* is the number of standard measurements with valid predictions from every assigned fit in bin *h*, *K_i_* is the number of calibration fits predicting observation *i*, 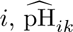 is its pH predicted by fit *k*, and pH*_i_* is its measured pH. Each acid-standard observation was predicted only by the fit that withheld its preparation, whereas each Fe_2_(SO_4_)_3_-mixture observation was predicted by all fits. The prediction root-mean-square error (RMSE; pH units) was calculated using:

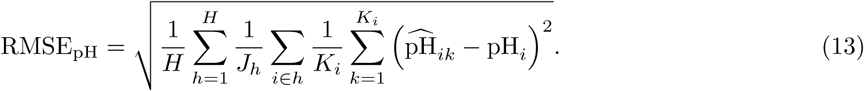

The propagated pH SD was calculated from the within-standard SD of the indicator and the local inverse-calibration slope^52^:

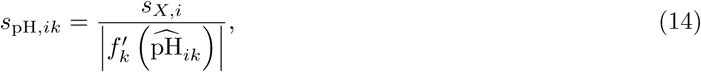

where *s_X,i_* is the within-standard sample SD of the indicator and 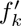 is the derivative of the fitted response in Equation (3), evaluated at the predicted pH. Measured and scatter-corrected predictions were paired by microplate and standard preparation. Measurements from each well were resampled together, with replacement within each microplate and standard preparation, and the models were refitted for each resample.^53^ Percentile confidence intervals^53^ were calculated from the refitted models. Intervals across microplates include between-preparation variation only where the microplates received independently prepared standard series; otherwise, they describe variation among technical microplates receiving the same preparation.

For absorbance comparisons between Fe_2_(SO_4_)_3_ and FeSO_4_ standard series at 380 and 428 nm, the FeSO_4_ absorbance was interpolated between adjacent measured total Fe concentrations to the matched concentrations in the Fe_2_(SO_4_)_3_ standard series. Absorbance was averaged over these concentrations within each microplate, retaining the pairing between wavelengths. Mean differences and Welch confidence intervals were calculated between the separately made Fe_2_(SO_4_)_3_ and FeSO_4_ standard series, each represented by technical microplates receiving the same preparation; these intervals describe technical microplate variation. Fe^3+^ calibration curves were fitted separately for each H_2_SO_4_ concentration using an empirical Weibull-type function:

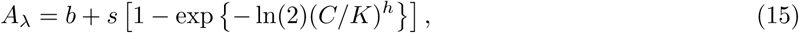

where *C* is the measured total Fe concentration (M), *b* is the fitted absorbance at zero Fe concentration (AU), *s* is the fitted response span (AU), *K* is the half-response concentration (M), and *h* is dimensionless. Fits included zero-Fe measurements, with *b* unconstrained and *s*, *K*, and *h* positive. Parameter and fitted-response intervals used the microplate-level procedure described for the pH calibrations. Shared and H_2_SO_4_-concentration-specific fits were compared by withholding each microplate or each intermediate concentration in the standard series across all microplates.^51^ Absorbance prediction errors and inverse-concentration bias and RMSE were calculated from predictions within the fitted absorbance or concentration range; the proportion of predictions excluded by these criteria was calculated. Relative concentration errors excluded the zero-Fe standard. Overlapping validation fits were not treated as independent replicates for confidence intervals or hypothesis tests.^54^

Mean Fe^2+^ concentrations determined by cerimetry were regressed against measured total Fe concentrations within each FeSO_4_ or Fe_2_(SO_4_)_3_ standard series by ordinary least squares. Slopes were compared with zero and unity and intercepts with zero within the comparison family comprising these tests across the FeSO_4_ standard series. Uncertainty in the total-Fe concentrations was not propagated into the regressions. Confidence intervals and hypothesis tests were calculated only when the residual variance was greater than zero. Apparent Fe^3+^ concentration in well *i* ([Fe^3+^]_app_*_.,i_*; M) was calculated using:

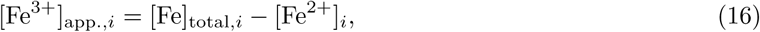

where [Fe]_total_,*_i_* and [Fe^2+^]*_i_* are concentrations calculated on the final-well basis from measurements of the corresponding standard: total Fe by ICP-OES after serial dilution and Fe^2+^ by cerimetry, respectively (M). Differences between conditions with and without dye, and between scatter-corrected and measured absorbance, were averaged over matched standards within each microplate and compared with zero across microplates. The comparison family comprised the dye effect on measured and scatter-corrected *A*_380_, and the scatter-correction effect with and without dye.

For handling tests, lid, microplate-material, temperature, and membrane effects were calculated as paired differences at matched standards. These differences were expressed as percentages of the comparator’s observed response span calculated using Equation (8), and averaged within each microplate. Temperature comparisons used the initial acquisition for the same material, microplate-material comparisons used matched standard preparations, and membrane comparisons used the preceding overnight acquisition without membrane. Paired microplate means were compared with zero within separate comparison families for lid effects, pH-indicator handling effects, and Fe^3+^-indicator handling effects. Changes in observed response span and pooled within-standard SD were calculated on matched complete wells and standards using Equation (11), with handling and comparator conditions as the numerator and denominator, respectively, and summarized across microplate pairs. To quantify separation between adjacent standards relative to well-to-well variability, the signed adjacent-standard separation score (*D_j_*; unitless) was calculated using:

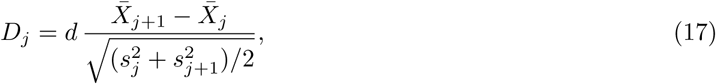

where *d* = *−*1 for the ratiometric pH indicator and *d* = 1 for the Fe^3+^ absorbance indicator. Positive scores indicate separation in the expected response direction, whereas negative scores indicate a reversal. The median adjacent-standard score was calculated for each microplate at each acquisition. Only originally adjacent standards with eligible paired technical-well measurements, estimable within-standard SDs, and a finite, positive denominator were included.

To assess the effects of extended microplate incubation on the indicators, the change from the initial measurement for well *i* at time *t* (Δ*X_i,t_*; AU or unitless) was calculated using:

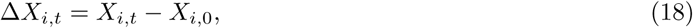

where *X* is 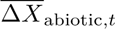 (AU) or *R*(∗) (unitless) and *X_i,_*_0_ is its initial measurement. Wells were averaged within each microplate and standard before calculating changes as percentages of the initial observed response span (*P_j,t_*; %):

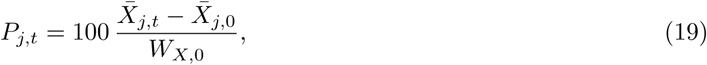

where subscripts *t* and 0 denote the acquisition time and initial acquisition, respectively, and *W_X,_*_0_ is the initial observed span calculated using Equation (8). Storage log-fold changes were calculated using Equation (11), with *X̅_j,t_* and *X̅_j,_*_0_ as the numerator and denominator, respectively. Log ratios were averaged over standards within each microplate and then across microplates. Storage effects were summarized at the final acquisition and by the mean, maximum absolute, and root-mean-square changes over time. Final-acquisition percentage changes and log ratios were compared with zero across microplates within a comparison family spanning the indicators and dye conditions. A sensitivity analysis retained only standards with valid measured and scatter-corrected measurements at every acquisition, using their original initial observed response spans. The availability of measured and scatter-corrected ratios was summarized within each microplate before equal-weight averaging across microplates. Concentration-matched edge–interior differences in Δ*X* were expressed as percentages of the initial observed response span (*P*_edge_,*_j,t_*; %) using:

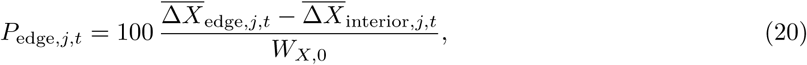

where the numerator is the difference between the mean changes of edge and interior wells at the same standard, calculated using Equation (18). Values from Equation (20) were averaged over matched standards within each microplate at each acquisition to obtain *P̅*_edge_,*_t_*. These differences were summarized at the final acquisition and by the time-weighted mean (*P*_edge_,_time_; %):

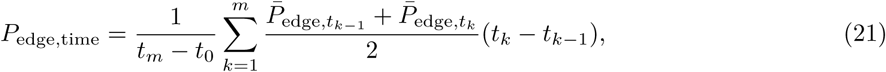

where *t*_0_*, …, t_m_* are the ordered acquisition times. These paired differences were compared with zero within the comparison family covering final-acquisition and time-averaged edge–interior differences across indicators and conditions.

Handling heatmaps used Equation (11) to express absorbance or derived ratios relative to a common comparator configuration. For pH, the numerator was the measured or scatter-corrected mean ratio for each standard within a microplate. The denominator was the matched measured mean ratio from the initial UV-transparent acquisition for temperature comparisons or the overnight UV-transparent acquisition without membrane for membrane comparisons. Log ratios were averaged over standards within each microplate and then across microplates. For Fe, the reference was initial measured *A*_380_ in UV-transparent microplates without dye at 23 *^◦^*C in 10^0^ M H_2_SO_4_. Absorbance was averaged across microplates for each standard before calculating log ratios. The numerator was the mean for the comparison condition, and the denominator was the reference mean absorbance interpolated between adjacent concentrations to the same total-Fe concentration. These log ratios were averaged over standards.

For medium compatibility, pooled standard deviations, observed response span, and *Q* used standards with estimable within-standard SDs. Calculations were performed separately for each medium and indicator, with arithmetic means across microplates. For eligible positive means, approximate confidence limits used the delta method on the logarithm of the mean^52^ and were back-transformed using:

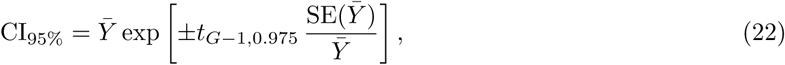

where *Y̅* is the arithmetic mean across microplates, SE(*Y̅*) is its standard error, and *G* is the number of eligible technical microplates receiving the same medium preparation. Adjacent-standard differences were calculated using Equation (5). Fe concentrations in these comparisons specify the added amount above the Fe already present in the medium, calculated from the measured standard concentration and the mixing volumes. The separate paired comparison of *Q* used Equation (11) on exactly the same eligible wells for both absorbance indicators, with scatter-corrected and measured values as the numerator and denominator, respectively. In this comparison using the same wells, all eligible standard means contributed to the observed response span; the pooled within-standard variance used only standards with estimable within-standard SDs. Tests against zero used technical microplate estimates within the comparison family of estimable medium-specific scatter-correction effects. Medium-background differences between conditions with and without dye were evaluated separately at 485, 545, and 750 nm, with the comparison family spanning these wavelengths and media.

For the biological demonstration tests, changes from initial measurements were calculated separately for each biotic and abiotic well using Equation (18). Technical wells were averaged within each microplate and biotic or abiotic condition to obtain the independent microplate estimates used in comparisons and figures; well-level values also supported technical-precision summaries. The net biotic change for well *i* at time *t* (Δ*X*_net_,*_i,t_*; AU or unitless) was calculated using:

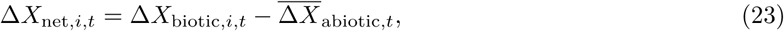

where Δ*X*_biotic_,*_i,t_* is the change in the biotic well and 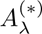 is the arithmetic mean change of condition-matched abiotic wells within the same microplate. Primary microplate–shake-flask comparisons used biotic Δ*A*_380_ as the microplate indicator of apparent Fe^3+^ generation^39^ and apparent Fe^3+^ generation in shake flasks, without abiotic subtraction. Apparent Fe^3+^ generation in shake flasks ([Fe^3+^]_app_*_.,f,t_*; M) was calculated using:

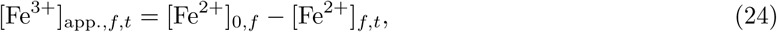

where [Fe^2+^]_0*,f*_ is the concentration measured in the shared starting medium and assigned to shake flask *f*, and [Fe^2+^]*_f,t_* is the concentration determined by cerimetry in its sample collected at time *t* (M).

Each time course was summarized by area under the curve (AUC) and time to the first observed maximum.^55^ AUC was calculated by signed trapezoidal integration from the initial acquisition to the first observed maximum using:

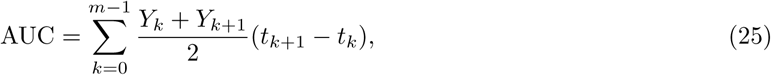

where *Y_k_* is the value of the measured or derived quantity at acquisition time *t_k_* (d), *t*_0_ is the time of the initial acquisition, and *m* is the index of the first observed maximum. For measurements made on withdrawn culture samples, *t_k_*denotes sample-collection time. AUC has units of the integrated quantity multiplied by days: AU*·*d for absorbance, d for a dimensionless ratio, OD_600_*·*d for apparent planktonic cell density expressed as OD_600_, and M d for apparent Fe^3+^ generation. Negative responses were retained, and tied maxima were assigned to the earliest acquisition. Integration required finite measurements spanning a nonzero duration and no missing measurements between the initial acquisition and the first observed maximum. Time to first observed maximum (*t*_max_; d) was measured from the initial acquisition using:

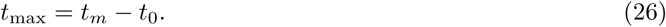

Trajectories with no positive rise had zero AUC at the initial maximum and were excluded from timing comparisons. Microplate and shake-flask summaries were calculated separately for each control or treatment condition, averaged within each experimental batch, and then averaged equally across batches.^56^ Between-batch variation was expressed as the sample coefficient of variation (CV; %) using:

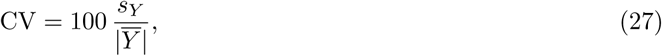

where *Y* and *s_Y_* are the arithmetic mean and sample standard deviation of batch means for the same condition, respectively. The deviation of batch *b* from this mean (*B_b_*; %) was calculated using:

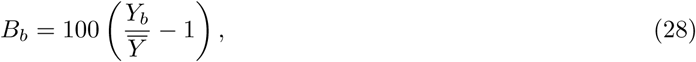

where *Y_b_*is the batch mean AUC or time to first observed maximum calculated using Equation (25) or Equation (26), respectively. Between-batch variances of natural-log AUC and time to first observed maximum were compared between no-additive AFM1 control conditions in microplates and shake flasks using two-sided *F* tests, assuming independent, normally distributed logarithmic values. Variance-ratio confidence intervals were calculated from the *F* distribution; the comparison family comprised the planned tests for AUC and time to first observed maximum. Exploratory shake-flask comparisons among Ga_2_(SO_4_)_3_ treatment conditions at 0, 10*^−^*^3^, and 10*^−^*^1^ M on a cation basis paired the same experimental batches and used two-sided paired Pitman–Morgan tests on natural logarithms of batch-level AUC before normalization to controls and time to first observed maximum.^57^ The effect was the ratio of the variances of log outcomes; marginal 95% CIs were obtained by inversion of the paired test. The comparison family comprised all planned condition-pair comparisons for AUC and time to first observed maximum. The assumption that the paired logarithmic values jointly follow a normal distribution could not be assessed adequately, so these comparisons were exploratory.

The screening criterion was biotic Δ*A*_380_ *≥* 1 AU and condition-matched abiotic Δ*A*_380_ *<* 1 AU at the same subsequent acquisition. A microplate met the criterion if an acquisition satisfied both thresholds. Classification as not meeting the criterion required valid initial measurements and complete paired subsequent measurements. Otherwise, the microplate outcome could not be evaluated; missing measurements were not interpreted as absence of apparent Fe^2+^-oxidation activity. Net biotic changes and differences from the thresholds were calculated for the same paired acquisitions, with each measurement counted once. Classification consistency was described by the range of microplate outcomes within each experimental batch and the range of fractions meeting the criterion between batches for the same condition. These comparisons required every constituent microplate classification to be evaluable. Treatment comparisons used assigned control conditions for the same measured or derived quantity, cultivation format, and experimental batch. AUC as a percentage of the same-batch control mean (*P*_AUC_; %) was calculated using:

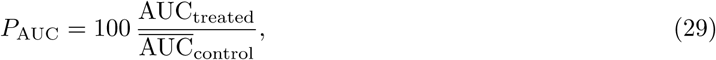

where the numerator is the treated-culture AUC and the denominator is the arithmetic mean control AUC, with each culture integrated separately to its first observed maximum using Equation (25). All assigned controls were required, and their mean AUC had to be positive. The difference in time to first observed maximum (Δ*t*_max_; d) was calculated using:

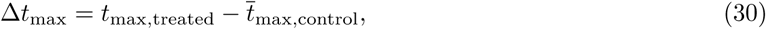

where *t*_max_,_treated_ is the treated-culture time to first observed maximum calculated using Equation (26) and *t*_max_,_control_ is the arithmetic mean for the assigned controls. Timing comparisons required a positive increase in the treated culture and every assigned control. Repeated treatment effects were described by between-batch standard deviations, agreement in direction, and Spearman rank correlations between batches using eligible shared conditions with nonconstant responses. Correlations were averaged across batch pairs within each additive and experimental preparation before calculating the median across these groups. Sensitivity to acquisition timing and observation duration was assessed by removing the final acquisition, retaining the first 75% of the observation duration, retaining alternate acquisitions plus the initial and final acquisitions, or treating the middle acquisition as missing. Classifications, AUCs, and timing were recalculated with the original missing observations and eligibility criteria retained. Screening consistency and sensitivity to acquisition timing and observation duration were summarized descriptively.

For shake flasks and culture tubes, apparent planktonic cell density expressed as OD_600_ was calculated from the fluorescence measurements as previously described using the calibration for *A. ferrooxidans*.^4^ Shake-flask estimates additionally used subtraction of the mean abiotic-control value of 0.001 in OD_600_ units. Negative, flagged, or missing fluorescence measurements were excluded from conversion. Measured and scatter-corrected microplate *A*_600_ were paired within wells. Ga_2_(SO_4_)_3_-associated AUC as a percentage of the same-batch control mean and difference in time to first observed maximum were calculated separately for each measurement type using Equation (29) and Equation (30), respectively. Confidence intervals were calculated across independent experimental batches where eligible; otherwise, comparisons were summarized by the available between-batch or within-batch standard deviation.

Citric acid effects at the final acquisition were calculated relative to zero-citric-acid control conditions at each Ga_2_(SO_4_)_3_ concentration (cation basis). Microplate comparisons paired treatments and controls within the same microplate, final acquisition, and elapsed time. Shake-flask comparisons paired cultures from the same experimental batch and inoculum at the same Ga_2_(SO_4_)_3_ concentration and final sample-collection time. Effects on net biotic change and on the matched abiotic measurements were calculated separately for microplates. Paired differences were averaged within experimental batch, and confidence intervals were calculated across paired experimental batches. For wavelength comparisons, the difference in net absorbance change (*W_p_*; AU) was calculated within microplate *p* at the final acquisition using:

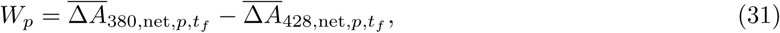

where 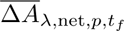 is the microplate mean net absorbance change at wavelength *λ* and final acquisition *t_f_*, calculated using Equation (23). These differences were averaged within experimental batch and compared with zero within the comparison family of citric acid/Ga_2_(SO_4_)_3_ formulations. The descriptive wavelength comparison summarized final net absorbance changes separately for each wavelength and treatment condition using equally weighted experimental-batch means and their sample SD.

Microplate–shake-flask correspondence in mean within-format AUC as a percentage of the same-batch control mean and mean time to first observed maximum was described using Equation (11), with microplate and shake-flask values as the numerator and denominator, respectively. The quantities were calculated using Equation (29) and Equation (26), respectively, distinguishing equal from neighboring tested concentrations. For trajectory-shape comparisons, net measured and scatter-corrected microplate *A*_380_ and fractional Fe^2+^ depletion in shake flasks were restricted to their shared observation period. Each trajectory was shifted to zero at the start of this period and divided by its positive change from start to end using:

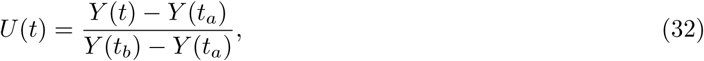

where *U* (*t*) is the trajectory scaled to its change over the shared observation period (unitless), *Y* (*t*) is the measured or derived quantity, and *t_a_*and *t_b_*are the start and end of that period (d). The time-weighted mean absolute difference (MAE*_t_*; unitless) between the scaled trajectories was calculated using:

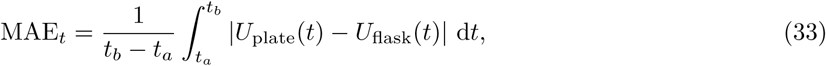

The time-weighted root-mean-square difference (RMSE*_t_*; unitless) was calculated using:

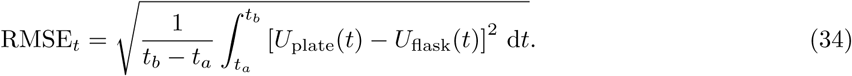

Integrals in Equation (33) and Equation (34) were evaluated over piecewise-linear trajectories, including zero crossings when integrating absolute differences. Comparisons of measured and scatter-corrected trajectories used the same eligible cultures and required eligible measurements throughout the shared observation period. Repeated comparisons were averaged within each shake-flask condition and microplate experimental batch before calculating an equally weighted mean across these combinations. The influence of integration duration was assessed separately by recalculating primary AUCs as percentages of the same-batch control mean over a shared time window.^55^ This window extended from the latest initial acquisition to the earliest final acquisition across both formats and all assigned controls, including observations after the maximum. The original primary measurements, cultures, controls, and eligibility criteria were retained; window boundaries were interpolated between adjacent finite observations. Changes in AUC as a percentage of the same-batch control mean and the microplate–shake-flask difference were summarized descriptively.

For the *L. ferriphilum* microplate tests, net changes were calculated at the final acquisition and summarized with the sample SD across independently inoculated microplates per condition within each experiment. The contextual *A. ferrooxidans* reference was interpolated between adjacent finite observations to the *L. ferriphilum* final-acquisition time and summarized across experimental batches as described above. For culture-tube validation experiments using fresh starter cultures, final biotic–abiotic differences (*D_Y,_*_final_) were calculated using:

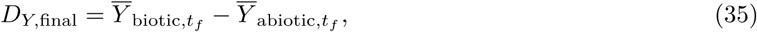

where *Y* _biotic_,*_t_* and *Y* _abiotic_,*_t_* are the arithmetic means of the measured variable *Y* for condition-matched biotic and abiotic culture tubes at the final acquisition time *t_f_*, with one contribution per culture tube. *D_Y,_*_final_ has the units of the measured variable. These differences used the final measurements rather than changes from measurements of the shared starting medium. The fluorescence conversion described above was not validated for *L. ferriphilum*.

For soluble-S microplate experiments, net changes in measured and scatter-corrected ratiometric pH indicators were calculated at the final acquisition. Scatter-correction effects were compared within wells with eligible paired final measurements using net indicator change standardized by abiotic measurement SD (*Z_i,t_*; unitless):

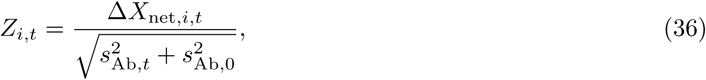

where *s*_Ab_,*_t_* and *s*_Ab_,_0_ are the sample standard deviations of condition-matched abiotic wells at the final and initial acquisitions, respectively. This score describes the net change relative to abiotic measurement SD rather than a hypothesis-test statistic. Signed scores and absolute scores were calculated for individual wells and averaged separately within each microplate before calculating scatter-corrected-minus-measured differences. The availability of measured and scatter-corrected measurements and the direction of paired responses were also compared. S_4_O_6_^2−^ culture-tube final pH and fluorescence measurements were calculated using Equation (35). Comparisons restricted to an individual experimental batch, or involving reused starter cultures, were summarized descriptively without treating repeated use of a starter culture as additional independent replication.

### Use of generative artificial intelligence

Models from ChatGPT (OpenAI), including GPT-5.6 Sol Ultra and GPT-6/6.2 Astra Ultra, were used to develop the structure of the data deposit, develop and implement the Data processing codes, develop and implement visuals, and verify consistency within and across the deposited dataset and this manuscript. The authors reviewed and executed the code, compared the plotted values with the source data, and validated its outputs.

### Data Records

The Zenodo deposit contains measurements and experimental documentation organized into six folders within the data/ directory, including experiments not displayed in the Technical Validation figures.^41^ The 00_materials and strains folder describes chemicals, materials, equipment, strains, medium formulations, and experimental conditions in Markdown tables and machine-readable YAML files. The 01 indicator calibration folder contains absorbance spectra, selected-wavelength measurements, storage and configuration comparisons, and measured standard pH values and Fe concentrations. The 02 media validation folder contains abiotic absorbance measurements grouped by indicator and medium, measured pH values of medium–standard mixtures, and the measured total Fe concentration of the calibration stock. The 03 biological microplates folder contains absorbance time courses organized by organism and treatment condition, including condition-matched abiotic control conditions. Its chronology subfolder contains acquisitions.csv and condition timecourse index.csv, which link reader exports to acquisition times, elapsed times, microplates, and experimental conditions. Acquisitions are ordered by timestamp, with elapsed time measured from the first deposited acquisition of each microplate. The 04 flask validation folder contains flask measurements.csv, with one row per shake flask and sample-collection time, including pH, oxidation–reduction potential (ORP), cerimetry, soluble elemental concentrations, and (Non-)planktonic fluorescence where available. The 05 test tube validation folder contains test tube measurements.csv, with culture-tube pH or ORP time courses and final planktonic fluorescence measurements where available. Its metadata also describe the qualitative follow-up using the original *L. ferriphilum* microplate inocula. Accompanying YAML files link measurements to experimental batches, vessels, wells, treatment and control conditions, and inocula, identifying repeated acquisitions and shared starting-medium measurements assigned to multiple vessels. Microplate-reader files with the .xls extension contain UTF-16LE tab-delimited text; acid-iron.xlsx is an Excel workbook, and measurement tables use UTF-8 CSV format. Unavailable or inapplicable values use NA in CSV files and null in YAML files, distinct from numerical zero. Instrument flags and finite negative measurements are retained. Accompanying descriptions and analysis status fields distinguish missing measurements, undefined calculations, exclusions, and results unavailable because of insufficient replication. The README.md files describe organization, variables, units, concentration bases, and time definitions. The repository contains analysis code in analyses/ and Figures 1–8, Tables 1–6, and Supplementary Tables S1–S32 in visualizations/. Each main-text table links its experimental components to the underlying records. Each supplementary table presents its caption, column definitions, and numerical results in one Markdown file.

**Figure 1:**
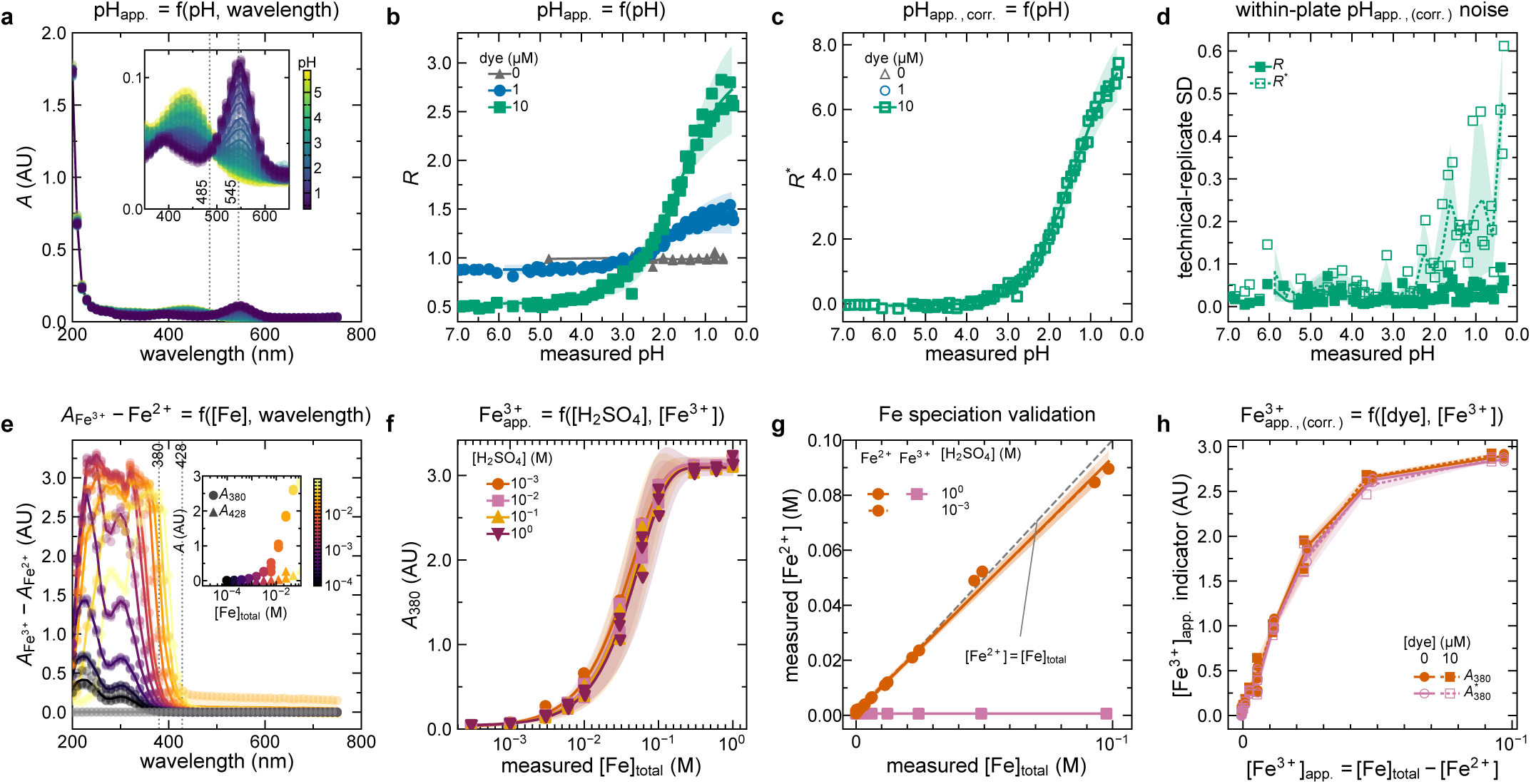
Ratiometric pH and Fe^3+^ absorbance indicators in chemical standards. (a) Spectra of H_2_SO_4_ standards containing nominally 10*^−^*^5^ M thymol blue, identified by measured pH; inset lines mark 485 and 545 nm. (b) Measured *R* = *A*_545_*/A*_485_ versus pH. (c) Scatter-corrected 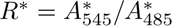 versus pH. Gray triangles, blue circles, and green squares identify nominal 0, 10*^−^*^6^, and 10*^−^*^5^ M thymol blue, respectively; filled and open symbols denote measured and scatter-corrected ratios. Curves are four-parameter logistic fits, except for measured ratios without dye, where lines connect means. (d) Within-standard technical-well standard deviation (SD) at 10*^−^*^5^ M thymol blue; solid and dashed lines denote *R* and *R^∗^*, respectively. (e) Difference between Fe_2_(SO_4_)_3_ and FeSO_4_ spectra interpolated to matched measured total Fe concentrations in 10^0^ M H_2_SO_4_ without dye; the inset compares the selected-wavelength measurements at 380 and 428 nm. (f) *A*_380_ versus measured total Fe concentration at different nominal H_2_SO_4_ concentrations, with separate nonlinear fits; zero-Fe standards are omitted from the logarithmic axis. (g) Mean Fe^2+^ concentration from replicate titrations of each standard versus measured total Fe concentration, with ordinary least-squares fits and a dashed identity line. (h) Measured *A*_380_ and scatter-corrected 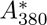 versus apparent Fe^3+^ concentration with or without dye. Optical points show one estimate per physical microplate and standard, after averaging eligible wells or, in (d), calculating their SD. Points in (b–d) use each preparation’s measured pH. The fits use mean measured pH for each standard. In (e), each point subtracts the interpolated mean ferrous reference from one ferric microplate. The band combines ferric and ferrous measurement SDs. Unfitted lines and bands indicate means *±* SD; fitted bands indicate 95% confidence intervals.

**Figure 2:**
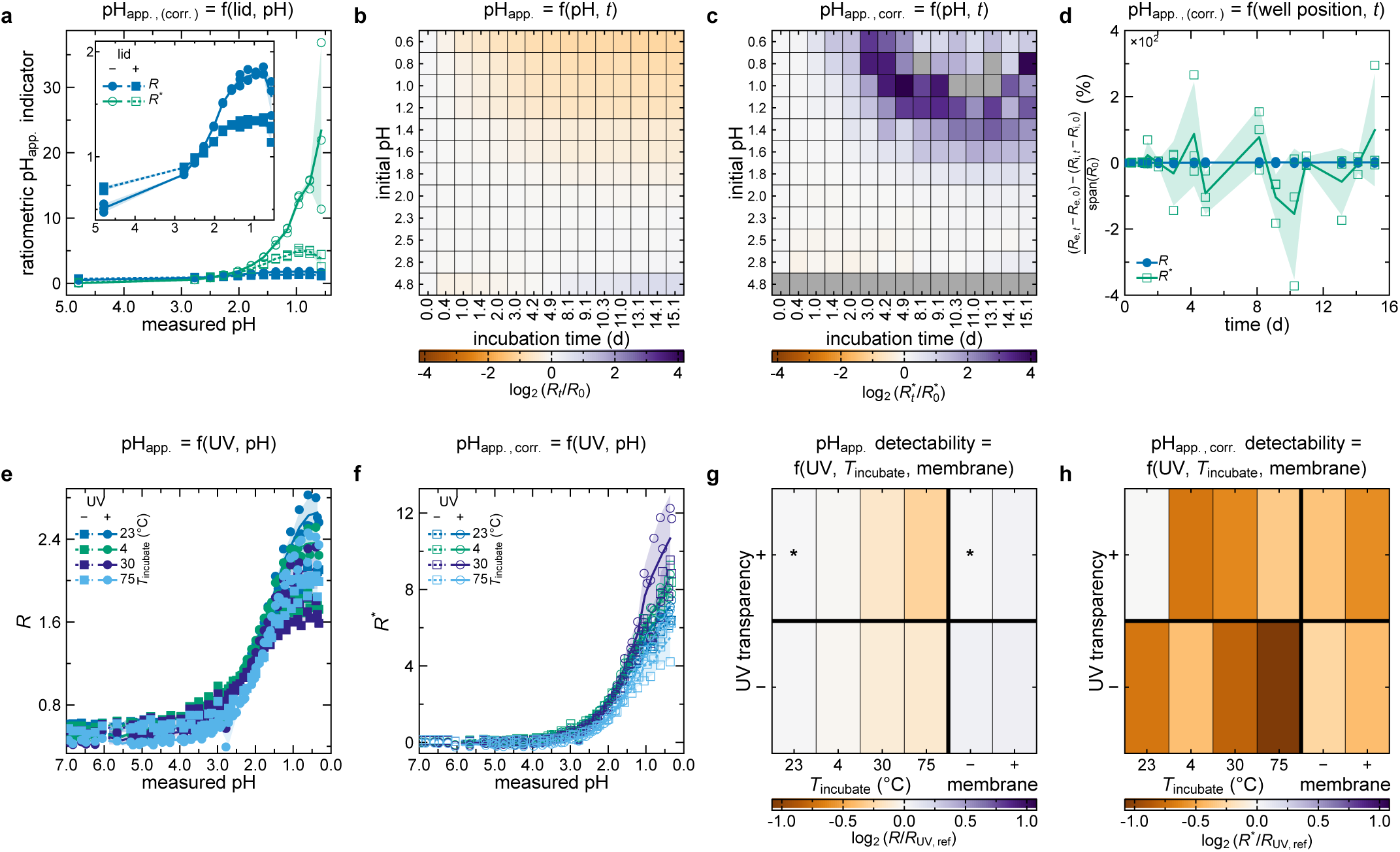
Ratiometric pH indicator measurements across microplate configurations and incubation conditions. (a) Measured thymol blue ratio *R* = *A*_545_*/A*_485_ and scatter-corrected *R^∗^* versus measured pH in UV-transparent microplates with or without a lid; the inset enlarges the lower-response region. (b) Mean log_2_(*R_t_/R*_0_) during benchtop incubation in non-UV-transparent microplates. (c) Mean 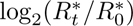 under the same storage conditions. (d) Edge–interior difference in the change from the initial indicator value, expressed as a percentage of the initial observed response span. (e) Measured *R* after sequential incubation at 23, 4, 30, and 75 *^◦^*C in UV-transparent and non-UV-transparent microplates. (f) Scatter-corrected *R^∗^* after the same sequential incubations. (g) Mean log_2_ ratio of measured *R* to the measured comparator *R*; asterisks identify the comparator conditions. (h) Mean log_2_ ratio of scatter-corrected *R^∗^* to the same measured comparator *R*. Temperature comparisons use the initial UV-transparent acquisition at 23 *^◦^*C; membrane comparisons use the overnight UV-transparent acquisition without membrane. Gray heatmap cells indicate unavailable ratios. In (a, d–f), each point represents one physical microplate. Lines and shading show the mean and between-microplate standard deviation at each standard or acquisition.

**Figure 3:**
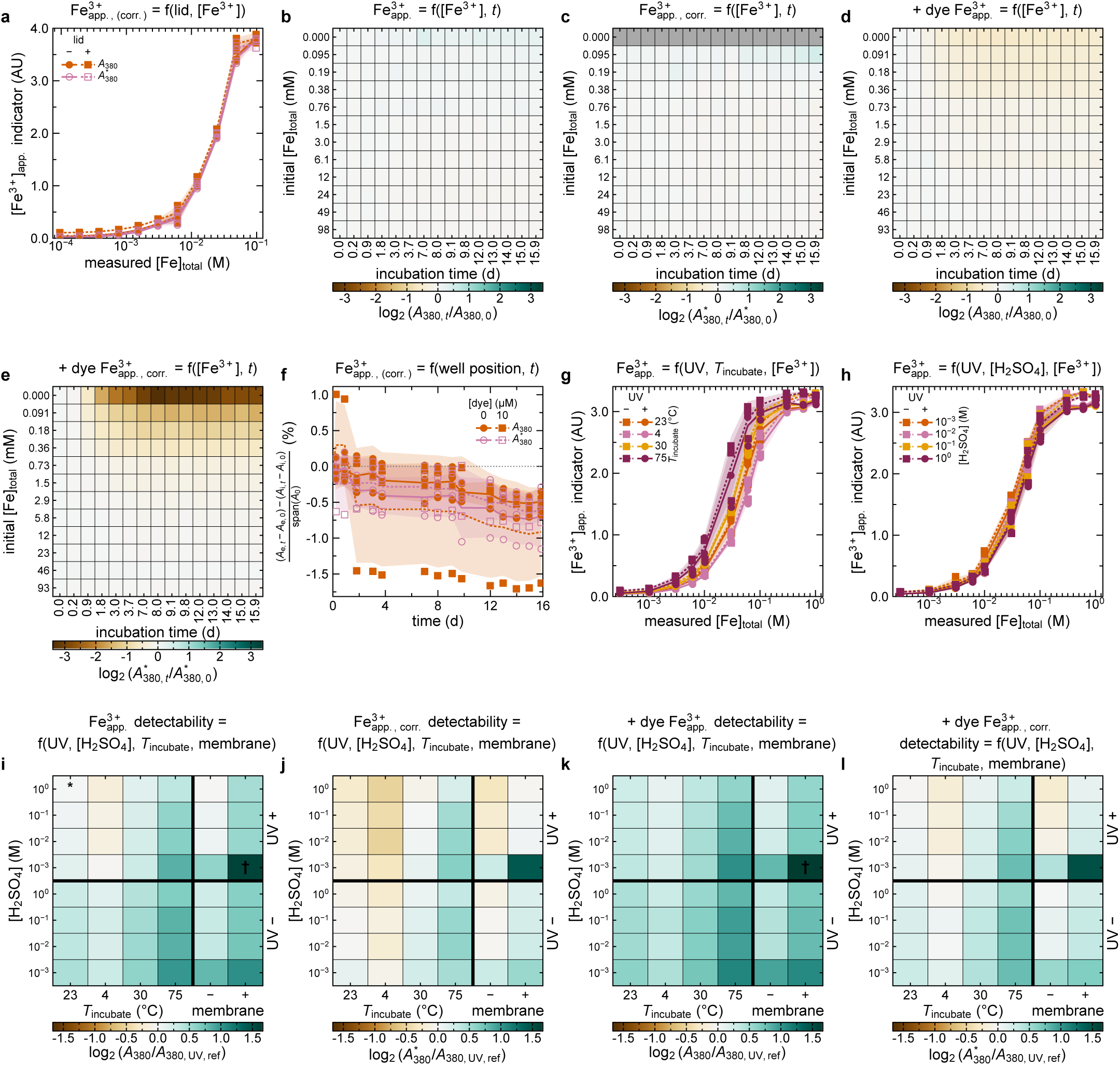
Fe^3+^ absorbance indicator measurements across microplate configurations and incubation conditions. (a) Measured *A*_380_ and scatter-corrected 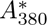 versus measured total Fe concentration with or without a lid. (b) Mean log_2_(*A*_380_,*_t_/A*_380_,_0_) during benchtop incubation without dye. (c) Mean 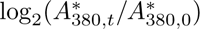 during benchtop incubation without dye. (d) Mean log_2_(*A*_380_,*_t_/A*_380_,_0_) during benchtop incubation with nominally 10*^−^*^5^ M thymol blue. (e) Mean 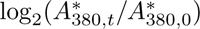 during benchtop incubation with nominally 10*^−^*^5^ M thymol blue. (f) Edge–interior difference in the change from initial 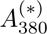, expressed as a percentage of the initial observed response span, for both dye conditions. (g) Measured *A*_380_ during sequential temperature incubations in 10^0^ M H_2_SO_4_ without dye. (h) Measured *A*_380_ versus measured total Fe concentration at different nominal H_2_SO_4_ concentrations in UV-transparent and non-UV-transparent microplates. (i) Mean log_2_ ratio of measured *A*_380_ without dye to the common comparator; the asterisk identifies the comparator condition. (j) Corresponding ratio for scatter-corrected 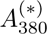 without dye. (k) Corresponding ratio for measured *A*_380_ with nominally 10*^−^*^5^ M thymol blue. (l) Corresponding ratio for scatter-corrected 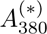 with nominally 10*^−^*^5^ M thymol blue. The common comparator is initial measured *A*_380_ in UV-transparent microplates without dye at 23 *^◦^*C in 10^0^ M H_2_SO_4_, interpolated to matched measured total Fe concentrations. Gray heatmap cells indicate unavailable ratios. Daggers identify the low-acid membrane acquisition with reversed concentration ordering. In (a, f–h), points represent physical microplates and lines and shading show their mean *±* standard deviation. Lid and storage comparisons use repeated microplates of shared preparations. Temperature and acid-background comparisons use independently prepared standards.

**Figure 4:**
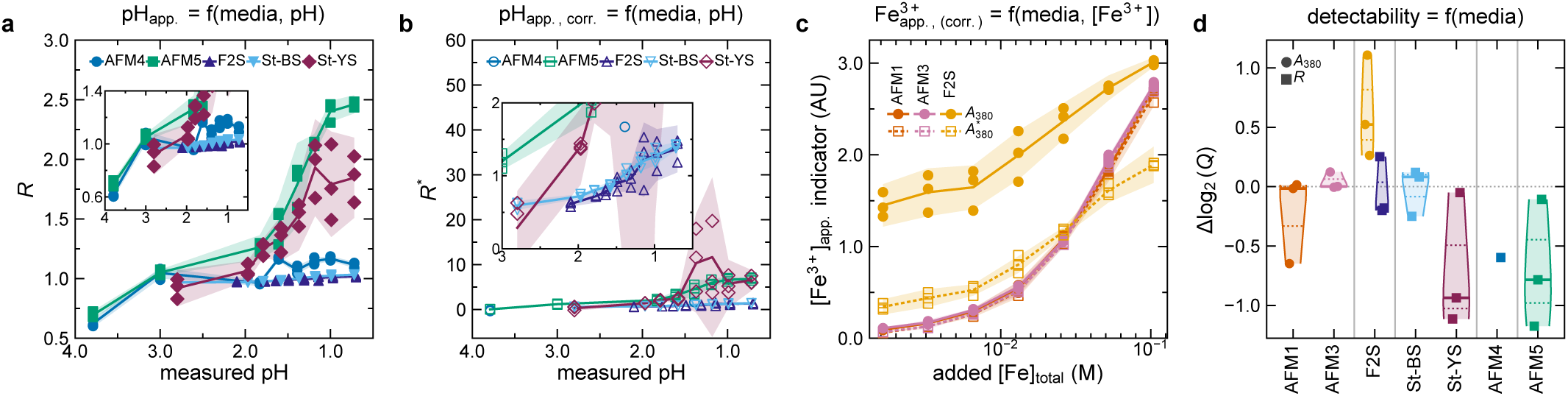
Ratiometric pH and Fe^3+^ absorbance indicators in acidophile media. (a) Measured thymol blue ratio *R* = *A*_545_*/A*_485_ versus measured pH in AFM4, AFM5, F2S, and Starkey (St) medium containing brown or yellow S^0^, abbreviated bS^0^ and yS^0^, respectively. (b) Scatter-corrected *R^∗^* versus measured pH in the same media; the insets enlarge the lower-response regions. (c) Measured *A*_380_ and scatter-corrected 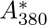 versus additional total Fe supplied as Fe_2_(SO_4_)_3_ to AFM1, AFM3, and F2S without dye; the horizontal axis excludes Fe already present in each medium. In (a–c), each point is the mean of eligible wells for one standard on one physical microplate. Lines show means across eligible wells and shading shows their pooled technical standard deviation (SD). Microplates sharing a standard preparation can give overlapping points. (d) Log_2_ ratio of the scatter-corrected to measured response-span-to-SD ratio, log_2_(*Q^∗^/Q*), calculated using the same eligible wells for each pair. Violin distributions, median and quartile marks, and individual microplate estimates summarize technical precision; AFM4 lacks sufficient eligible estimates for an across-microplate SD or hypothesis test. Vertical separators distinguish Fe^2+^ media (AFM1 and AFM3), medium containing Fe^2+^ and bS^0^ (F2S), bS^0^ and yS^0^ media (St), S_2_O_3_^2−^ medium (AFM4), and S_4_O_6_^2−^ medium (AFM5). In (d), circles denote *A*_380_ and squares denote *R*. F2S includes both indicators.

**Figure 5:**
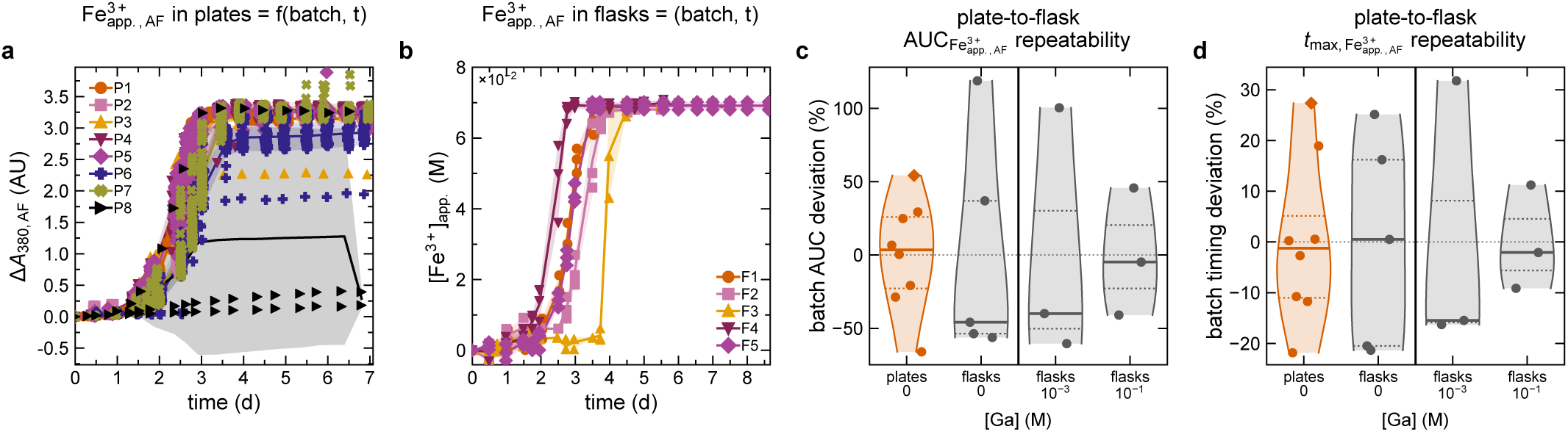
Between-batch variation in microplate and shake-flask measurements for *A. ferrooxidans*. (a) Biotic change from initial *A*_380_ in no-additive AFM1 microplate cultures across batches P1–P8, without abiotic subtraction. (b) Apparent Fe^3+^ generation estimated from Fe^2+^ depletion in no-additive AFM1 shake-flask cultures across batches F1–F5. Points represent independent microplate or shake-flask estimates; lines and shaded regions indicate within-batch means *±* standard deviation (SD). (c) Between-batch percentage deviations in area under the curve (AUC) for the corresponding indicator or apparent Fe^3+^ generation, including shake-flask treatments with 10*^−^*^3^ or 10*^−^*^1^ M Ga_2_(SO_4_)_3_ on a cation basis. (d) Corresponding percentage deviations in time to first observed maximum. AUC extends from the initial measurement to the first observed maximum of each trajectory. Violin distributions show median and quartile marks, with each point representing an experimental batch. Diamonds identify the grouped batch containing no-additive control conditions from iodate experiments. A vertical separator distinguishes no-additive control conditions from Ga_2_(SO_4_)_3_ treatments.

**Figure 6:**
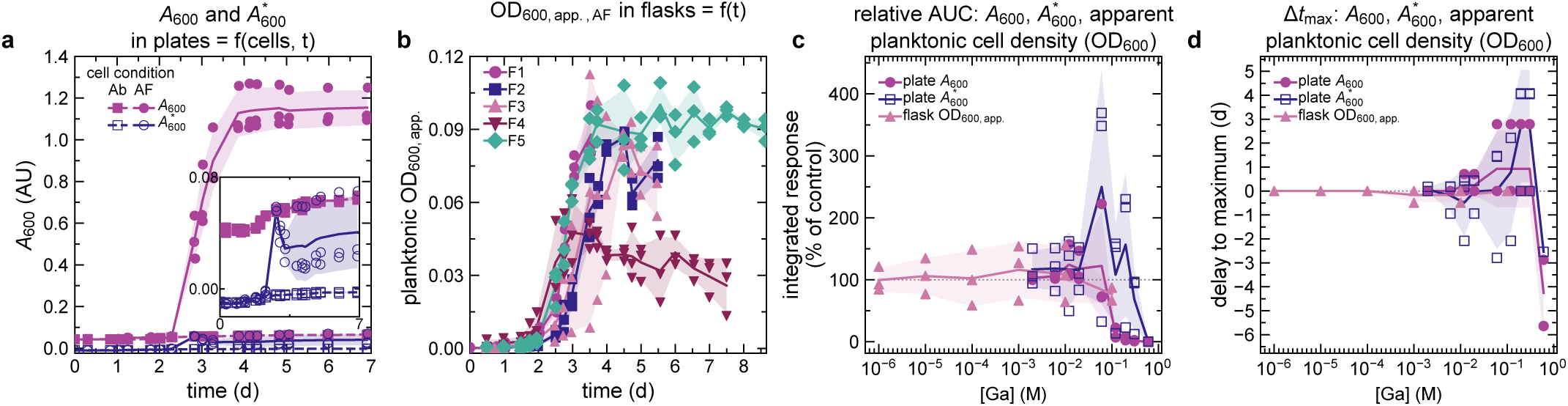
Microplate absorbance and apparent planktonic cell density in shake flasks for *A. ferrooxidans*. (a) Measured *A*_600_ and scatter-corrected 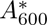 for paired no-additive *A. ferrooxidans* (AF) and abiotic (Ab) conditions; open symbols denote scatter correction. The inset enlarges both Ab series and the scatter-corrected AF series. (b) Fluorescence-derived apparent planktonic cell density expressed as OD_600_ in no-additive shake-flask cultures, after abiotic-control subtraction. Points represent independent microplate or shake-flask estimates; lines and shaded regions indicate within-batch means *±* standard deviation (SD). (c) Area under the curve (AUC) as a percentage of the same-batch, same-measurement no-additive control mean, versus Ga_2_(SO_4_)_3_ treatment concentration on a cation basis. (d) Treatment-minus-control differences in time to first observed maximum for the same measurement types. Concentration-response lines connect within-batch means at the respective nominal concentrations or concentrations calculated from measured stocks; shaded regions indicate within-batch vessel SD. The no-additive control is omitted from the logarithmic concentration axis.

**Figure 7:**
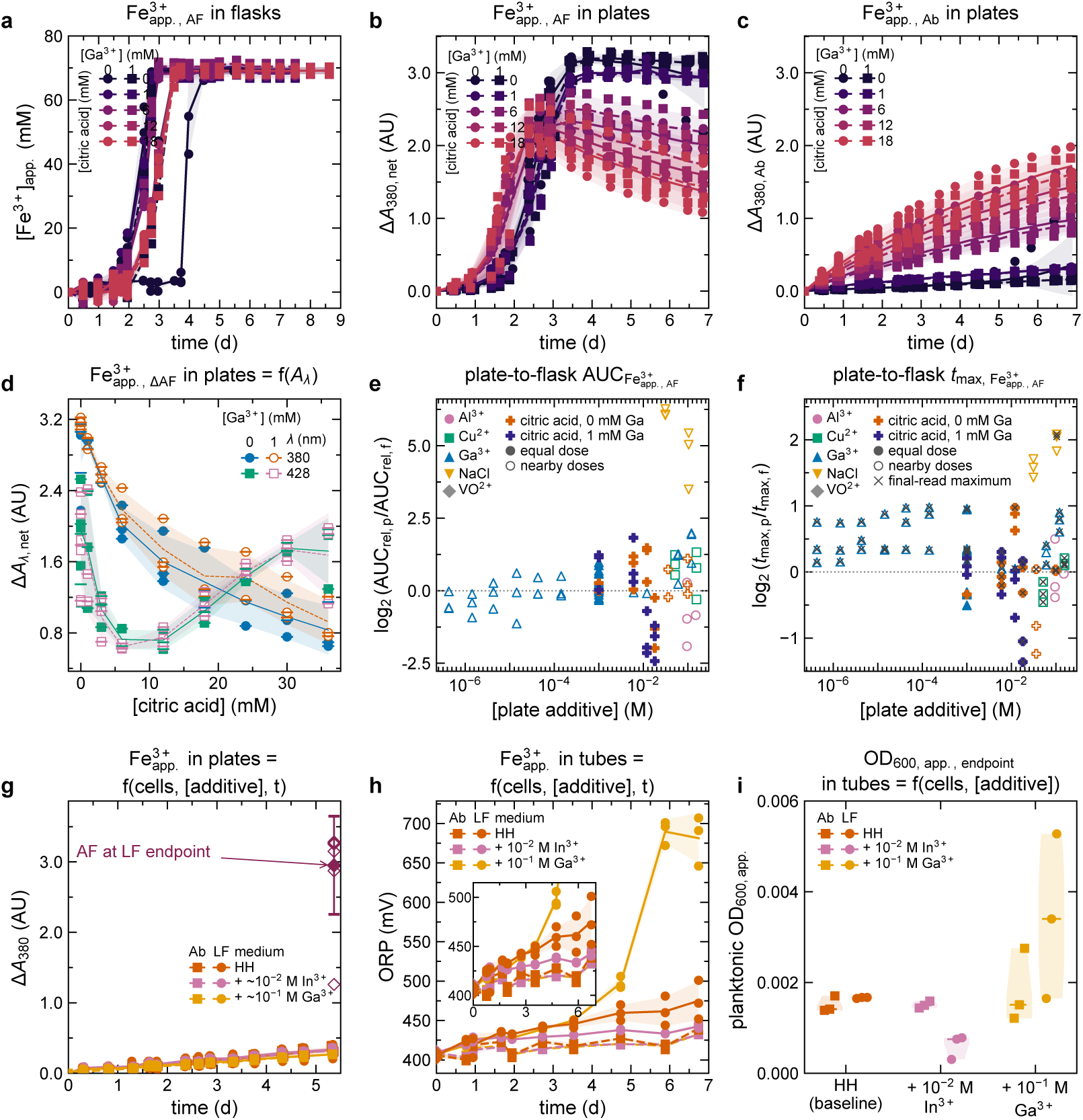
Microplate and shake-flask comparisons and *L. ferriphilum* validation measurements. (a) Shake-flask apparent Fe^3+^ generation estimated from Fe^2+^ depletion for *A. ferrooxidans* (AF) citric-acid/Ga_2_(SO_4_)_3_ conditions. (b) Net biotic microplate Δ*A*_380_ for the same treatment formulations. (c) Corresponding abiotic (Ab) Δ*A*_380_. In (a–c), color identifies citric acid concentration; circles/solid lines and squares/dashed lines denote 0 and 1 mM added Ga^3+^, respectively. Net change subtracts mean condition-matched abiotic change from biotic change since the initial measurement. (d) Net changes at the final acquisition at 380 nm (circles) and 428 nm (squares) versus citric acid; filled/open symbols and solid/dashed lines denote 0 or 1 mM added Ga^3+^, respectively. Points show paired microplates. Horizontal ticks show batch means. Lines and bands show the equally weighted mean *±* between-batch standard deviation (SD). (e) Log_2_ microplate/shake-flask ratio of relative area under the curve (AUC), log_2_(AUC_rel_,_p_*/*AUC_rel_,_f_), versus microplate additive concentration. (f) Log_2_ microplate/shake-flask ratio of mean time to first observed maximum, log_2_(*t̅*_max_,_p_*/t̅*_max_,_f_), versus microplate additive concentration. Relative AUC is the percentage of the same-batch no-additive control mean. Filled/open symbols denote equal/neighboring concentrations; crosses identify maxima at the final acquisition. Labels Al^3+^, Cu^2+^, Ga^3+^, and VO^2+^ denote added Al_2_(SO_4_)_3_, CuSO_4_, Ga_2_(SO_4_)_3_, and VOSO_4_, respectively; citric acid series contain 0 or 1 mM added Ga^3+^. Metal-salt concentrations use a cation basis. (g) *L. ferriphilum* (LF) and Ab changes from initial *A*_380_ at 1.428 *×* 10*^−^*^1^ M added Ga^3+^ or 6.22 *×* 10*^−^*^3^ M added In^3+^, with HH controls; Open diamonds show the eight AF batch means interpolated at 5.366 d. The filled diamond and error bar show their mean *±* SD. (h) Culture-tube oxidation–reduction potential relative to Ag/AgCl with 3 *×* 10^0^ M KCl, using fresh LF starters; diamonds denote shared starting-medium measurements. (i) Fluorescence-derived apparent planktonic cell density expressed as OD_600_ using the AF conversion, for the fresh-starter culture tubes; violins show individual culture tubes and medians. In trajectories and final absorbance comparisons, points represent vessels and lines and bands show means *±* SD. For LF comparisons, orange, pink, and yellow denote HH, In_2_(SO_4_)_3_, and Ga_2_(SO_4_)_3_, respectively; circles/squares and solid/dashed lines distinguish LF/Ab.

**Figure 8:**
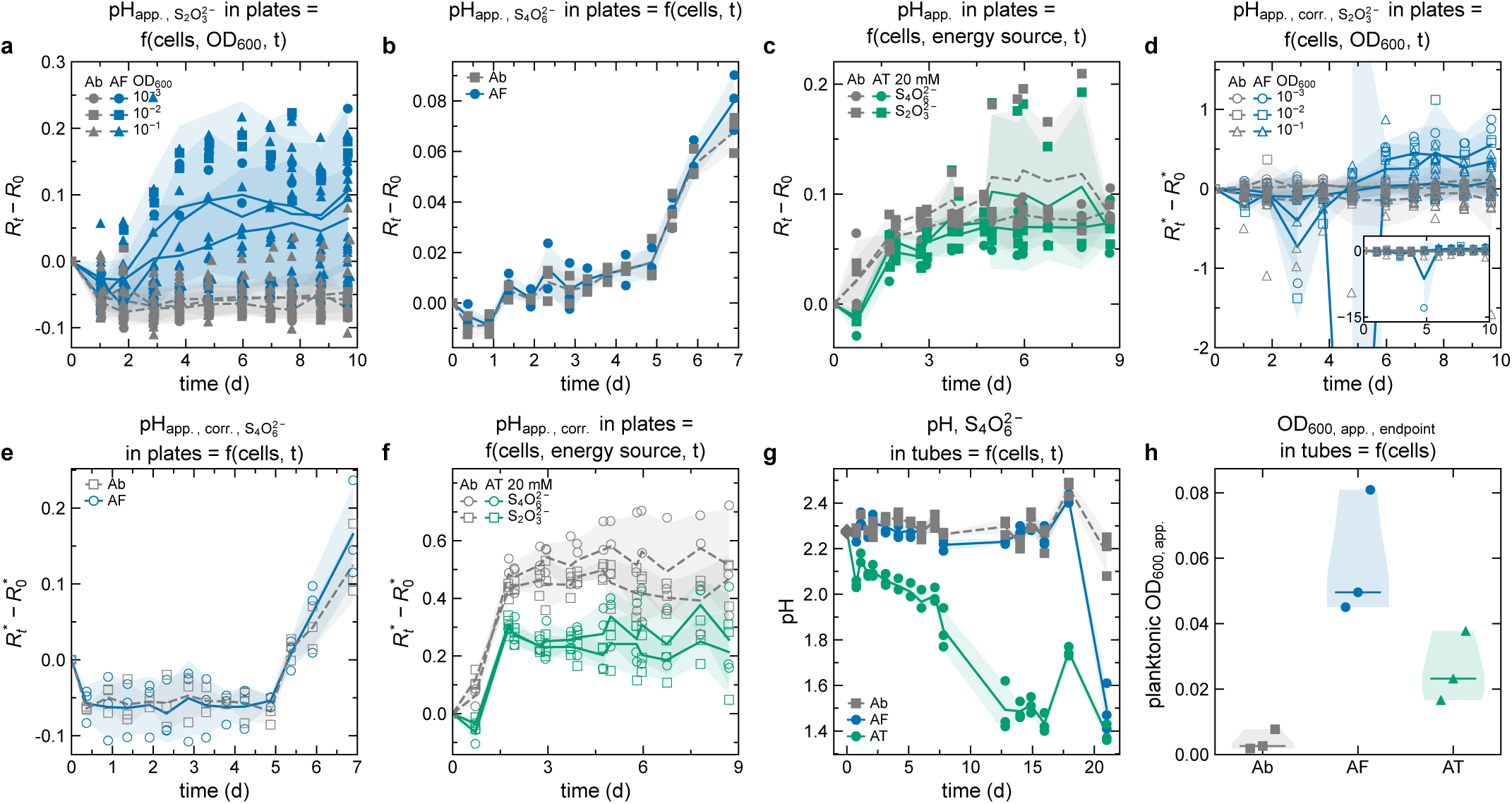
Ratiometric pH indicators in soluble-S microplate cultures and S_4_O_6_^2−^ culture-tube validation. (a) Changes from the initial measured thymol blue ratio *R* = *A*_545_*/A*_485_ for *A. ferrooxidans* (AF) supplied with S_2_O_3_^2−^. (b) Changes from initial measured *R* for AF supplied with S_4_O_6_^2−^. (c) Changes from initial measured *R* for *A. thiooxidans* (AT) supplied with S_2_O_3_^2−^ or S_4_O_6_^2−^. (d) Changes from initial scatter-corrected *R^∗^* for AF supplied with S_2_O_3_^2−^ ; the inset shows the full indicator range. (e) Changes from initial scatter-corrected *R^∗^* for AF supplied with S_4_O_6_^2−^. (f) Changes from initial scatter-corrected *R^∗^* for AT supplied with S_2_O_3_^2−^ or S_4_O_6_^2−^. Nominal substrate concentrations before dye addition were 2.015 *×* 10*^−^*^2^ M S_2_O_3_^2−^ for AF, 5 *×* 10*^−^*^3^ M S_4_O_6_^2−^ for AF, and 2 *×* 10*^−^*^2^ M of either substrate for AT; AT treatments contained 10*^−^*^5^ M nominal added H_2_SO_4_. Inoculum OD_600_ values also refer to the mixtures before dye addition; Ab denotes abiotic controls. Filled and open symbols denote measured and scatter-corrected ratios, respectively. Microplate points represent within-plate means; lines and shaded regions show means *±* standard deviation (SD) across eligible microplates, with gaps where the required measurements were unavailable. (g) Measured pH trajectories in TB-04 culture tubes containing 5 mM K_2_S_4_O_6_ without added Ga_2_(SO_4_)_3_; circles and squares show individual tubes, diamonds show each shared starting-medium measurement once, and lines and bands show means *±* SD. (h) Apparent planktonic cell density expressed as OD_600_ at the final culture-tube acquisition, using the AF-derived fluorescence conversion; violins show individual measurements and median marks. Gray, blue, and green identify Ab, AF, and AT, respectively.

### Technical Validation

Feasibility tests examined the concentration ranges of H^+^ and/or Fe^3+^ over which the indicators could distinguish changes in pH or total Fe concentration (Tables 1–2). Thymol blue produced absorbance spectra that changed with the measured pH of H_2_SO_4_ standards (Figure 1a).^44^ The measured ratiometric pH indicator *R* generally decreased with increasing pH (Figure 1b). This behavior was describable by a four-parameter logistic model and generally monotonic, although some adjacent standards had the opposite ordering. Within the comparison family of three scatter-correction effects at 10*^−^*^5^ M thymol blue, scatter correction notably increased the observed response span (Holm-adjusted *p* = 0.00726; Figure 1c). It also notably increased the pooled within-standard SD (Holm-adjusted *p* = 0.00726; Figure 1d),^4,45^ though the change in the response-span-to-SD ratio was not statistically supported after Holm adjustment (*p* = 0.0505). At 0 and 10*^−^*^6^ M thymol blue, scatter-corrected ratios did not provide complete replicate groups for fitting or estimating within-standard SD. Scatter correction reduced pH prediction RMSE for withheld H_2_SO_4_ standards, but prediction errors were larger when these calibrations were applied to Fe_2_(SO_4_)_3_ mixtures over the pH range represented in both experiments. Separately, the difference between absorbance spectra of Fe_2_(SO_4_)_3_ and FeSO_4_ standards was greater at 380 nm^39^ than at 428 nm^42^ (Figure 1e). *A*_380_ was therefore selected as the primary Fe^3+^ indicator. In Fe_2_(SO_4_)_3_ standards, *A*_380_ increased with concentration and approached a plateau at approximately 5 *×* 10*^−^*^2^–5 *×* 10*^−^*^1^ M Fe_2_(SO_4_)_3_ (Figure 1f). Separate empirical calibration curves described each H_2_SO_4_ concentration, but did not consistently improve prediction of withheld measurements. Inverse concentration predictions became less reliable near saturation, and adjacent-standard separation varied across the tested concentration range. Within the comparison family of six estimable regression tests across the FeSO_4_ standard series, slopes relating Fe^2+^ determined by cerimetry to measured total Fe were below unity in 10*^−^*^3^ M H_2_SO_4_ (Holm-adjusted *p* = 0.00769) and in 10^0^ M H_2_SO_4_ (Holm-adjusted *p* = 0.0117; Figure 1g). The Fe_2_(SO_4_)_3_ series had a constant cerimetry-equivalent response, including the zero-Fe standard. Consequently, total Fe minus the Fe^2+^ result was interpreted as the apparent Fe^3+^ concentration. Below saturation, measured and scatter-corrected *A*_380_ generally increased with apparent Fe^3+^ concentration (Figure 1h). Within the comparison family of four dye and scatter-correction contrasts, the with-versus-without-dye difference was not statistically supported for measured *A*_380_ (Holm-adjusted *p* = 1.00) or scatter-corrected 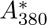 (Holm-adjusted *p* = 1.00). Together, these simulant measurements supported the use of thymol blue and *A*_380_ to detect acidification and apparent Fe^3+^ formation, respectively, as expected.^39,44^

Extendability tests measured changes in abiotic indicator responses during storage and under different acquisition and microplate configuration conditions. Within the comparison family of four lid-effect contrasts, the measured ratiometric pH indicator was lower with a lid than without one (Holm-adjusted *p* = 0.00487), as was its scatter-corrected counterpart (Holm-adjusted *p* = 0.0318; Figure 2a). In contrast, the lid effect was not statistically supported for measured *A*_380_ (Holm-adjusted *p* = 0.140) or scatter-corrected 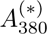 (Holm-adjusted *p* = 0.193; Figure 3a). Although indicator changes during benchtop storage did not differ significantly between edge and interior wells, edge wells were excluded from subsequent biological culture experiments as a precaution against potential edge effects. Scatter-corrected ratios more often failed the eligibility criteria, and deviations remained when comparisons included only standards with valid measurements throughout storage. Mean final changes were small relative to the initial observed spans for measured and scatter-corrected *A*_380_ without dye (Figure 3b–c). This was also true for the measured and scatter-corrected *A*_380_ with the dye (Figure 3d–e), although some standard concentrations showed larger changes during storage. Within the comparison family of 12 final-acquisition and time-averaged edge–interior contrasts, none was statistically supported after Holm adjustment (adjusted *p* = 0.238–1.00; Figure 2d; Figure 3f). Thus, these tests did not detect a systematic edge–interior difference in the measured indicator changes under the tested storage conditions. The pH indicators remained measurable during sequential temperature incubations in both microplate materials without and with scatter correction (Figure 2e–f). The Fe^3+^ indicator also remained measurable in sequential temperature comparisons (Figure 3g) and comparisons across nominal H_2_SO_4_ concentrations (Figure 3h) in both microplate materials. However, a similar observed response span did not imply a similar pooled within-standard SD or signed adjacent-standard separation score. After addition of gas-permeable membranes, responses remained measurable without and with scatter-correction for the ratiometric pH indicator (Figure 2g–h). This was also true for the Fe^3+^ indicator with (Figure 3i–j) and without dye (Figure 3k–l). Since the indicator responses remained broadly measurable and the non-UV-transparent plates were about 10*×* cheaper, we therefore limited further tests to the non-UV-transparent microplate configuration. These simulant measurements showed that the indicators remained detectable across multiple microplate configurations and incubation conditions, with generally monotonic responses and empirical calibration fits in the evaluated standard series, supporting a generalizable utility for the assay under wider acidophile cultivation conditions.

Compatibility tests examined whether medium composition altered the relationship between indicator response and measured pH or added Fe concentration (Table 3). The measured ratiometric pH indicator changed with measured pH in several media (Figure 4a). However, scatter correction did not consistently increase the observed response span or eliminate non-monotonic behavior between adjacent standards (Figure 4b). The response-span-to-SD ratio depended on medium composition. Smaller values were observed in yellow-S^0^ medium and in medium containing Fe^2+^ and brown S^0^ than in brown-S^0^ and S_4_O_6_^2−^ media. In S_2_O_3_^2−^ medium, scatter correction left too few standards with eligible replicate measurements to estimate the observed span under the primary eligibility criterion. In the Fe_2_(SO_4_)_3_-addition experiments, measured and scatter-corrected *A*_380_ increased monotonically between adjacent standards in Fe^2+^ media with and without citric acid (Figure 4c). The indicator response in medium containing Fe^2+^ and brown S^0^ had larger within-standard SDs and some reversed ordering (Figure 4c). Within the comparison family of seven estimable medium-specific contrasts, changes in the response-span-to-SD ratio after scatter correction were not statistically supported after Holm adjustment (*p ≥* 0.891; Figure 4d). AFM4 lacked sufficient eligible microplate estimates for an SD across microplates or a hypothesis test. The abiotic compatibility tests showed that these indicators could remain detectable and, in some media, monotonic despite non-target components, supporting medium-specific calibration of these absorbance indicators in the absence of bacteria.

Demonstration tests examined whether the same control and treatment conditions gave consistent microplate outcomes across experimental batches (Table 4). Across the Fe^2+^ treatment screen, most no-additive *A. ferrooxidans* control conditions satisfied the operational criterion for apparent Fe^2+^-oxidation activity, although satisfying that criterion did not necessarily imply a large net biotic change in *A*_380_. Classifications were generally consistent across experimental batches, as were the rime-course profiles from the baseline treatment condition (Figure 5a).

Validation tests examined whether microplate indicator changes corresponded to Fe^2+^ depletion, pH, ORP, and apparent planktonic cell density expressed as OD_600_ in shake flasks and culture tubes (Tables 5– 6). For no-additive AFM1 control conditions, the apparent oxidation of Fe^2+^ in microplates (Figure 5a) also increased in shake flask comparison tests (Figure 5b). Within the comparison family of two no-additive control-condition variance comparisons, between-batch variability did not differ detectably between the cultivation formats for log AUC (Holm-adjusted *p* = 0.617; Figure 5c) or log time to first observed maximum (Holm-adjusted *p* = 0.617; Figure 5d); however, these nonsignificant variance tests do not necessarily imply equal variability. Within the comparison family of six shake-flask treatment-pair and AUC/timing variance comparisons, none of the four estimable differences was statistically supported after Holm adjustment (adjusted *p* = 1.00 for each). The comparison of microplate *A*_600_ (Figure 6a) with fluorescence-derived apparent planktonic cell density expressed as OD_600_ in shake flasks (Figure 6b) examined whether the measurements gave consistent measured responses. Treatment AUC as a percentage of the same-batch control mean (Figure 6c) and differences in time to first observed maximum (Figure 6d) differed among measured *A*_600_, scatter-corrected *A*_600_, and apparent planktonic cell density. The *A*_600_ were therefore not considered to be a suitable proxy for the fluorescence-derived apparent planktonic cell density expressed as OD_600_ under the tested conditions. Apparent Fe^3+^ generation from shake flasks (Figure 7a) and microplates (Figure 7b) differed among citric-acid/Ga_2_(SO_4_)_3_ treatment conditions. The abiotic control conditions showed changes in *A*_380_ that depended on citric acid concentration (Figure 7c), indicating a confounding contribution from the medium chemistry. Within the comparison family of 18 citric-acid/Ga_2_(SO_4_)_3_ treatment formulations, none of the final differences between net Δ*A*_380_ and net Δ*A*_428_ remained statistically supported after Holm adjustment (adjusted *p* = 0.0845–0.628; Figure 7d). Agreement between cultivation formats varied among some treatment conditions for both AUC as a percentage of the same-batch control mean (Figure 7e) and time to first observed maximum (Figure 7f). Scatter correction did not consistently reduce differences between indicator trajectories and chemical measurements scaled to their changes over the shared observation period. Using the common observation interval instead of the time to first observed maximum also changed the treatment-to-control AUC comparisons. *L. ferriphilum* microplate cultures showed little separation in *A*_380_ from condition-matched abiotic control conditions (Figure 7g), in contrast to what was observed from non-identical *A. ferrooxidans* microplate cultures incubated for the same duration. Culture-tube follow-up with the original *L. ferriphilum* inocula did not show the visual change to a red color in HH medium used to assess apparent Fe^2+^-oxidation activity relative to abiotic controls through a 14 d observation period. Separate culture-tube experiments with fresh starter cultures showed positive final ORP differences relative to abiotic control conditions under some treatments (Figure 7h) and higher apparent planktonic cell density expressed as OD_600_ (Figure 7i). Together, these *L. ferriphilum* results could imply that the microplate cultures were not inoculated with viable cells. Small or variable biotic–abiotic differences in the measured ratiometric pH indicator occurred in *A. ferrooxidans* supplied with S_2_O_3_^2−^ (Figure 8a), *A. ferrooxidans* supplied with S_4_O_6_^2−^ (Figure 8b), and *A. thiooxidans* supplied with either soluble-S substrate (Figure 8c). Scatter correction changed the magnitude and sometimes the direction of these differences in *A. ferrooxidans* supplied with S_2_O_3_^2−^ (Figure 8d), *A. ferrooxidans* supplied with S_4_O_6_^2−^ (Figure 8e), and *A. thiooxidans* supplied with either substrate (Figure 8f), without consistently increasing net indicator change standardized by abiotic measurement SD. It also reduced the number of eligible ratios in some *A. ferrooxidans* treatment conditions. For (in)soluble-S experiments, direct pH and fluorescence measurements in culture tubes^4^ were used to examine the interpretation of small ratiometric pH indicator changes in the microplate experiments. In separate S_4_O_6_^2−^ culture-tube experiments, *A. ferrooxidans* and *A. thiooxidans* had lower final pH than abiotic control conditions (Figure 8g) and higher apparent planktonic cell density expressed as OD_600_ (Figure 8h). These results could imply that the microplate culture configuration was unsuccessful at generating distinguishable responses under some of the tested conditions, whereas differences could be resolved in the culture-tube configuration. Together, the biological comparisons supported microplate detection of apparent Fe^2+^ oxidation by viable *A. ferrooxidans* cultures, with more limited and condition-dependent support for acidification by soluble-S cultures.

## Usage Notes

The dataset supports assessment of indicator performance and comparisons among treatment conditions under the tested conditions. It may also support subsequent estimation of apparent kinetic parameters of metabolic byproduct formation and concentration–response relationships, although these biological models were not fitted here and their estimates will depend on model assumptions and observation periods.^42^ Users evaluating alternative corrections should begin with measured absorbances and apply the eligibility criteria in Data processing; values excluded from one calculation remain available for other appropriate analyses. Nominal concentrations, measured concentrations, and final concentrations calculated from additive stocks should remain distinguishable because precipitation, hydrolysis, complexation, and adsorption can alter soluble elemental concentrations or chemical availability.^58^

The abiotic-corrected absorbance at 380 nm indicates apparent generation of the metabolic byproduct Fe^3+^. The ratiometric pH indicator tracks changes in H^+^ activity, which can also arise from abiotic processes. Changes in apparent planktonic cell density expressed as *OD*_600_ do not have to track apparent Fe^2+^-oxidation activity.^36^ For cultures supplied with S_2_O_3_^2−^, S_4_O_6_^2−^, or brown S^0^, a biotic–abiotic pH-indicator difference is consistent with apparent oxidation of the specified substrate^4,23,40^ but does not quantify its conversion. The F2S compatibility tests characterize abiotic indicator responses in medium containing Fe^2+^ and brown S^0^. Medium composition, light scattering, precipitation, evaporation, speciation, and abiotic oxidation or reduction can affect the indicators; condition-matched abiotic control conditions and medium-specific calibration are therefore needed. Without suitable calibration, responses should remain expressed as indicator changes in their measured units. Inferring metabolic byproduct concentrations or equivalent substrate oxidation across media requires calibration. Complementary measurements are likely needed to assess biological interpretations and mechanisms (e.g., pH, Fe^2+^ determined by cerimetry, soluble Fe and sulfate concentrations, cell density, protein, DNA, microscopy).

Static microplate cultures differ from shake flasks and stirred or sparged systems in oxygen transfer, evaporation, gas exchange, surface-area-to-volume ratio, particle behavior, and mixing.^59^ These differences can affect oxidation of substrates requiring gas transfer or access to insoluble S,^23^ so treatment responses should be assessed in the cultivation format relevant to their intended application.

The assay may be applicable to chemolithotrophic acidophiles and other non-model microorganisms whose substrate oxidation produces a measurable change in these indicators. Extensions to additional species, consortia, engineered strains, media, or anaerobic conditions,^60^ require assessment under the intended chemical and cultivation conditions, using appropriate substrate-only, cell-free, and no-additive biotic control conditions. Compatibility with (gaseous) energy substrates other than those tested, including H_2_,^61^ would also need separate assessment.

## Data Availability

The experimental measurements, acquisition metadata, and source data for the figures and tables are available in the Zenodo dataset 10.5281/zenodo.24179050.^41^

## Author Contributions

**D.A.D.** and **M.A.H.** contributed equally to this work. The order of names was decided by mutual agreement among the these two authors.

**D.A.D.**: conceptualization (thymol blue as an indicator of acidification, experimental design), methodology (design across all five experimental series), investigation (indicator calibration, microplate cultures, and shake-flask measurements), validation, formal analysis, visualization, data curation, software (analysis and plotting code), data curation, writing – original draft (all unless noted otherwise), and writing – review & editing. **M.A.H.**: conceptualization (absorbance at 380 nm as an indicator of ferric iron formation), methodology (medium validation and ferric iron calibration), investigation (absorbance measurements and microplate culture experiments), validation, writing – original draft (methods), and writing – review & editing. **W.W.**: validation (assay performance across experimental formats), software (analysis and plotting code), data curation, writing – original draft (methods), and writing – review & editing. **L.N.M.**: methodology (design of soluble-sulfur culture experiments) investigation (microplate culture measurements and culture-tube endpoint fluorescence), and writing – review & editing. **S.T.B.**: methodology (design of soluble-sulfur culture experiments) investigation (culture-tube preparation and endpoint fluorescence measurements), and writing – review & editing. **S.B.**: writing – review & editing, resources, supervision, project administration, and funding acquisition.

## AI Use Declaration

The use of ChatGPT for code, manuscript, and figure preparation is described in Methods. We reviewed and verified the AI-assisted content and take responsibility for the final manuscript.

## Code Availability

All Python scripts used to analyze the data and generate the figures and tables are included in the Zenodo deposit and can also be accessed in the corresponding GitHub repository snapshot.^41^

